# In search of the Goldilocks zone for hybrid speciation II: hard times for hybrid speciation?

**DOI:** 10.1101/2023.02.15.528680

**Authors:** Alexandre Blanckaert, Vedanth Sriram, Claudia Bank

## Abstract

Hybridization opens a unique window for observing speciation mechanisms and is a potential engine of speciation. One controversially discussed outcome of hybridization is homoploid hybrid speciation by reciprocal sorting, where a hybrid population maintains a mixed combination of the parental genetic incompatibilities, preventing further gene exchange between the newly formed population and the two parental sources. Previous work showed that, for specific linkage architectures (i.e., the genomic location and order of hybrid incompatibilities), reciprocal sorting could reliably result in hybrid speciation. Yet, the sorting of incompatibilities creates a risk of population extinction. To understand how demographic consequences of the purging of incompatibilities interact with the formation of a hybrid species, we model an isolated hybrid population resulting from a single admixture event. We study how population size, linkage architecture and the strength of the incompatibility affect survival of the hybrid population, resolution/purging of the genetic incompatibilities and the probability of observing hybrid speciation. We demonstrate that the extinction risk is highest for intermediately strong hybrid incompatibilities. In addition, the linkage architecture displaying the highest hybrid speciation probabilities changes drastically with population size. Overall, this indicates that population dynamics can strongly affect the outcome of hybridization and the hybrid speciation probability.

## Introduction

The role of hybridization remains a central theme in the study of adaptation and speciation (Barton and Bengtsson, 1986; Buerkle et al., 2000; Barton, 2001; Mallet, 2007; Servedio et al., 2013; Schumer et al., 2018; Butlin et al., 2021). Hybridization plays an instrumental role in adaptive introgression (Hedrick, 2013; Seehausen, 2013; Meier et al., 2019) and promotes speciation through the subsequent increase in genetic variability (Marques et al., 2019). However, it can also increase the amount of genetic admixture to the point of reversing the speciation process (Seehausen et al., 2008) or lead to genetic or demographic population swamping (Todesco et al., 2016). These different consequences highlight the need to understand better the role of hybridization from both an evolutionary and ecological point of view.

Hybrid speciation is an additional outcome of hybridization; it refers to the formation of a new species upon admixture between two incipient species that subsequently become reproductively isolated from both parent species (Buerkle et al., 2000; Runemark et al., 2019). We distinguish between allopolyploid and homoploid hybrid speciation depending on whether or not the ploidy of the newly formed population differs from the parental populations. Contrary to the allopolyploid case, where the change in ploidy between the hybrid species and the parental species can immediately lead to reproductive isolation, reproductive isolation between homoploid hybrid and parental populations cannot be formed in a single step but rather has to evolve over time. For homoploid hybrid speciation to be possible, the parental populations must be genetically isolated, but the extent of this isolation must allow for hybridization events to still occur. Despite this apparent contradiction, several cases of homoploid hybrid speciation have been reported (Mavárez et al., 2006; Yakimowski and Rieseberg, 2014; Lamichhaney et al., 2018), and the matter has been extensively debated in the literature (Nieto Feliner et al., 2017; Schumer et al., 2018).

The *a priori* contradictory effects of reproductive isolation on hybrid speciation have been investigated through theoretical models (Buerkle et al., 2000; Schumer et al., 2015; Comeault, 2018; Blanckaert and Bank, 2018). Most of these models focus on the role of post-zygotic reproductive isolation. One classical model to explain the evolution of this type of genetic barrier is the classical Bateson-Dobzhansky-Muller model (Bateson, 1909; Dobzhansky, 1936; Muller, 1942). A (Batseon-)Dobzhansky Muller incompatibility (DMI) consists of two alleles at different loci whose mutual interactions are negatively epistatic and therefore deleterious when they occur in the same genome. A classical DMI can only isolate the hybrid population from a single parental source. As a consequence, a hybrid population can form its own reproductively isolated population only through the reciprocal sorting of multiple DMIs. Most previous models have focused on the population genetics of the newly formed hybrid population and its fate, assuming that the population sizes remained constant (with the exception of Buerkle et al. (2000), where population dynamics were included) and therefore ignoring the consequences of the genetic incompatibilities on the population dynamics of the hybrid population.

The role of population dynamics, and the choice to include or ignore them in evolutionary modeling, has been well studied and documented (for a review, see Ravigné et al. (2009)), with the two modeling regimes being named hard and soft selection, respectively. The terms were initially coined by Wallace (1968) and more formally defined by Christiansen (1975) (however these two regimes are not complementary (Wallace, 1975)). In the soft selection regime, selection acts locally on smaller subpopulations. Survival of a genotype is linked to the presence/absence of other genotypes, so soft selection is a frequency- and density-dependent process (Wallace, 1975). The number of adult individuals in each generation is fixed and always equals the population’s carrying capacity. Conversely, under a hard selection regime, selection operates at a global level, and the population size is variable, depending on the viability of individuals of the population. The viability is determined solely by the individual genotype and is both density and frequency independent (Wallace, 1975). While more complex than their soft selection counterparts, hard selection models capture both ecological and evolutionary dynamics of a population instead of just the evolutionary dynamics, allowing us in particular to investigate extinction events.

Here, we use a hard selection approach to determine whether conditions favorable to hybrid speciation are also favoring population extinction. Building upon previous soft selection models (Schumer et al., 2015; Blanckaert and Bank, 2018), we modeled both the demographics and evolutionary dynamics of a newly formed hybrid population following admixture. Reproductive isolation is caused by (intrinsic) post-zygotic genetic incomaptibilities (DMIs). We tracked the fate of the newly formed population, as well as the genes responsible for reproductive isolation. We found that survival of the population was strongly influenced by the linkage architecture (*i*.*e*., the order and distance of genes along the genome) and the initial population size of the hybrid population. Hybrid speciation was also possible under a hard selection regime. As seen previously, the linkage architecture had a large impact on the hybrid speciation probability (Blanckaert and Bank, 2018). However, which linkage architectures facilitate hybrid speciation and which architectures ensure higher survival was different and depended on the initial population size of the hybrid population. We also quantified the impact of both co-dominant and recessive epistasis and discussed the implications of these results on the study of hybrid speciation.

## Method

We model the evolution of a hybrid population in discrete, non-overlapping generations. We assume that the hybrid population is formed through a single admixture event of two parental populations with a proportional contribution of *i*_*p*_ and (1 *− i*_*p*_) (a hybrid swarm scenario), after which the hybrid population remains isolated. Initially, this newly formed population is assumed to have a total number of individuals of *N*_init_. Different from Blanckaert and Bank (2018), the size of the newly formed population evolves in response to selection and genetic drift.

We study evolution at four diallelic loci,*A*_1_, *A*_2_, *B*_1_ and *B*_2_, with the derived allele denoted by an uppercase letter and the ancestral by a lower-case letter. We assume that selection affected viability and model fitness of an individual as its probability to reach adulthood. All surviving individuals (*i*.*e*., adults) contribute equally to the gamete pool. Zygote formation is random. Therefore, ancestral alleles are under single-locus selection, with a fitness disadvantage *α*_*k*_ of allele *a*_*k*_ compared to *A*_*k*_. Similarly, allele *b*_*k*_ suffers from a fitness disadvantage *β*_*k*_ compared to allele *B*_*k*_. In addition, alleles *A*_*k*_ and *B*_*k*_ are epistatically interacting, forming a DMI, with a fitness cost *ϵ*_*k*_ associated with the interaction. Accordingly,

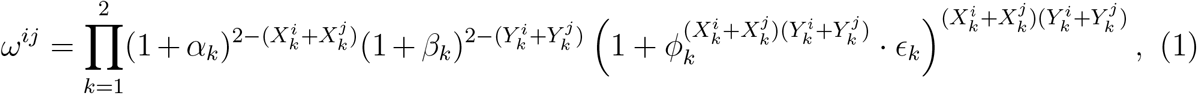

where 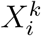 denotes the number of alleles *k* in haplotype *i* and 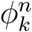 is a placeholder for the dominance of epistasis. If the DMI is codominant then *∀n ∈* {0, 1, 2, 4}, 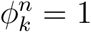; if the DMI is recessive, then *∀n ∈* {0, 2, 4}, 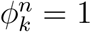, but for *n* = 1, 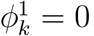.

Selection (hard selection) happens in the diploid phase of the life cycle and affects the survival probability of juveniles to adulthood, therefore impacting the number of adult individuals *N* (*t*) of the population at generation *t*. More precisely, the number of adult individuals of genotype *ij*, at generation *t* + 1, is drawn from a Poisson distribution, with parameter *λ*^*ij*^(*t*) = *ω*_off_ *\* ω*^*ij*^ *\* f* ^*ij*^(*t* + 1) *\* N* (*t*). The term *ω*_off_ corresponds to the mean number of offspring produced by the ancestral genotype (per default, *ω*_off_ = 1.01) and *ω*^*ij*^ to the fitness of genotype *ij* as defined above. The number of adult individuals at generation *t* is given by *N* (*t*) and *f* ^*ij*^(*t* + 1) represents the frequency of genotype *ij* in the juvenile pool (produced by the adults from generation *t*) assuming Hardy Weinberg equilibrium. The parameter *λ*^*ij*^(*t*) represents the expected number of surviving juveniles carrying genotype *ij* at generation *t* + 1. Finally, we assume that there is no density regulation of the population size; if the mean absolute fitness of the population is larger than 1, the population exponentially grows up to the carrying capacity of the environment, *K*, and is then maintained at this constant size.

The parental populations are assumed to be monomorphic for the following genotypes: *A*_1_*b*_1_*A*_2_*b*_2_*/A*_1_*b*_1_*A*_2_*b*_2_ and *a*_1_*B*_1_*a*_2_*B*_2_*/a*_1_*B*_1_*a*_2_*B*_2_, respectively. We are particularly interested in two specific long-term outcomes of the eco-evolutionary dynamics: extinction, and hybrid speciation. We define hybrid speciation as the reciprocal sorting of the genetic incompatibilities (Schumer et al., 2015; Blanckaert and Bank, 2018). More precisely, it corresponds to the fixation of a haplotype that is incompatible with both parental haplotypes, without carrying an incompatibility itself (i.e. either haplotype *A*_1_*b*_1_*a*_2_*B*_2_ or *a*_1_*B*_1_*A*_2_*b*_2_). For consistency, we always use the same nomenclature to refer to haplotypes, even if it does not match the physical order of the loci along the chromosome. There are a total of six possible linkage architectures (since *A* and *B* are interchangeable, see Fig. 1).

**Figure 1.**
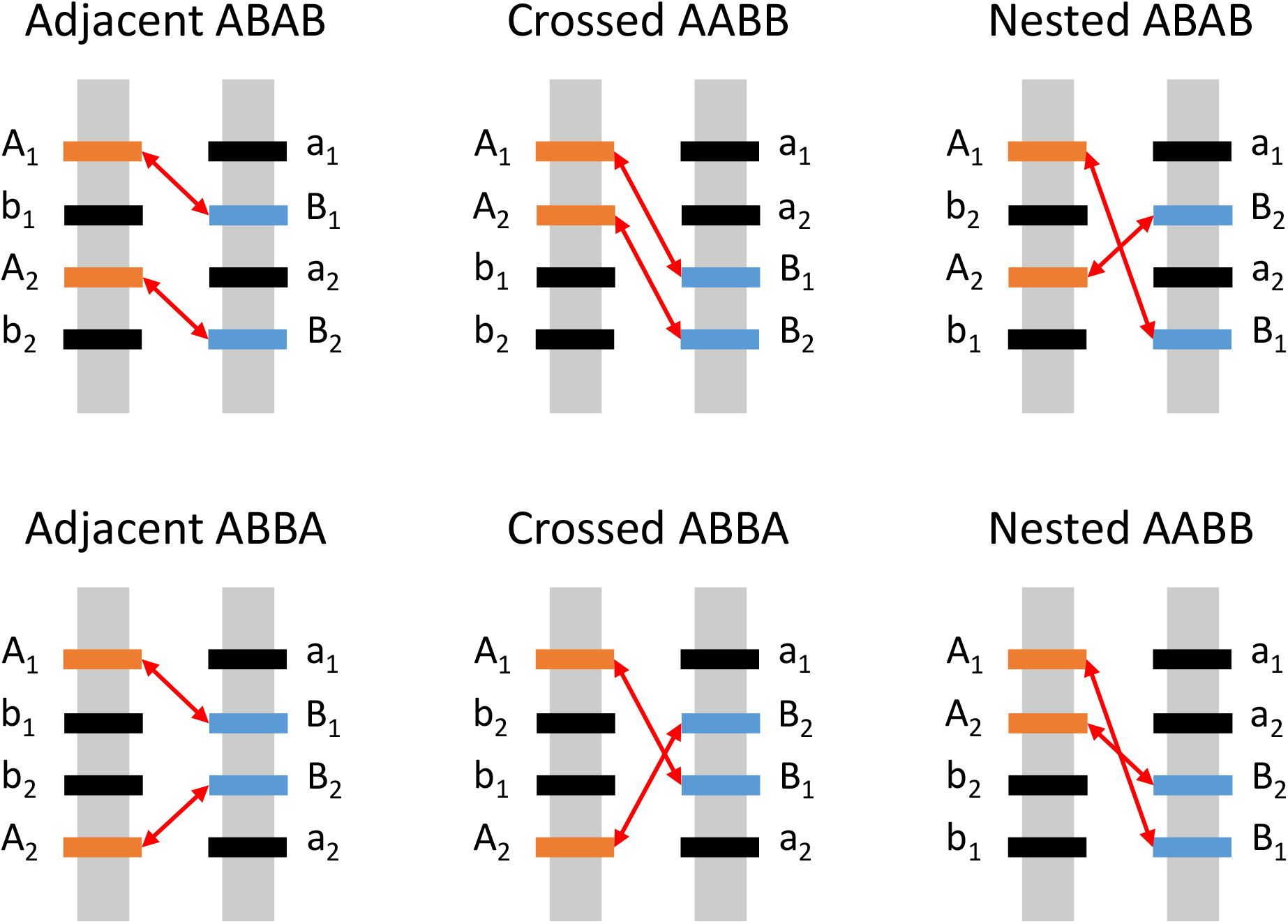
: All 6 possible linkage architectures. Each architecture is illustrated for an F1 hybrid individual. Derived alleles are given in orange (*A*_*k*_) and blue (*B*_*k*_); their ancestral counterparts are depicted in black. Red arrows indicate the epistatic interactions between derived alleles *A*_*k*_ and *B*_*k*_. This figure is adapted from Figure 1 in Blanckaert and Bank (2018), using the same naming convention.

Recombination between two adjacent loci is given by *r*_*XY*_ (0 *≤ r*_*XY*_ *≤* 0.5). Here, *r*_*XY*_ = 0.5 corresponds to a scenario in which the two loci are located on different chromosomes. Recombination between non adjacent loci *X* and *Y*, separated by a third locus *Z*, is given by *r*_*XY*_ = *r*_*XZ*_(1 *− r*_*ZY*_) + *r*_*ZY*_ (1 *− r*_*XZ*_). For more than three loci, the recombination rate can be calculated recursively.

When studying the case of two loci, the *A*_2_ and *B*_2_ loci are considered neutral (*α*_2_ = *β*_2_ = *ϵ*_2_ = 0) in our simulations, and their fate is ignored in the analysis.

## Results

### The hybrid population is at highest risk of extinction for intermediate values of epistasis or recombination

Compared to the soft selection regime, an additional evolutionary outcome arises when population sizes are dynamic: extinction of the newly formed hybrid population. First, we consider the case of unlinked loci (Fig. 2). The presence of genetic incompatibilities puts the newly formed hybrid population at risk of extinction, especially in small populations (see Fig. 2 and Fig. S1); the stronger the epistasis and the smaller the population, the higher the risk of extinction. This is true both for recessive and codominant incompatibilities (Fig. 2 A-B). A sufficiently large initial population size (*N*_init_ *>* 1, 000) can be sufficient to prevent extinction in the case of a recessive incompatibility (Fig. 2A). For a codominant DMI, even large populations (*N*_init_ = 10^4^) have an approximately 25% risk of extinction when the incompatibility is almost lethal (*ϵ* = *−*0.99; Fig. 2B).

**Figure 2.**
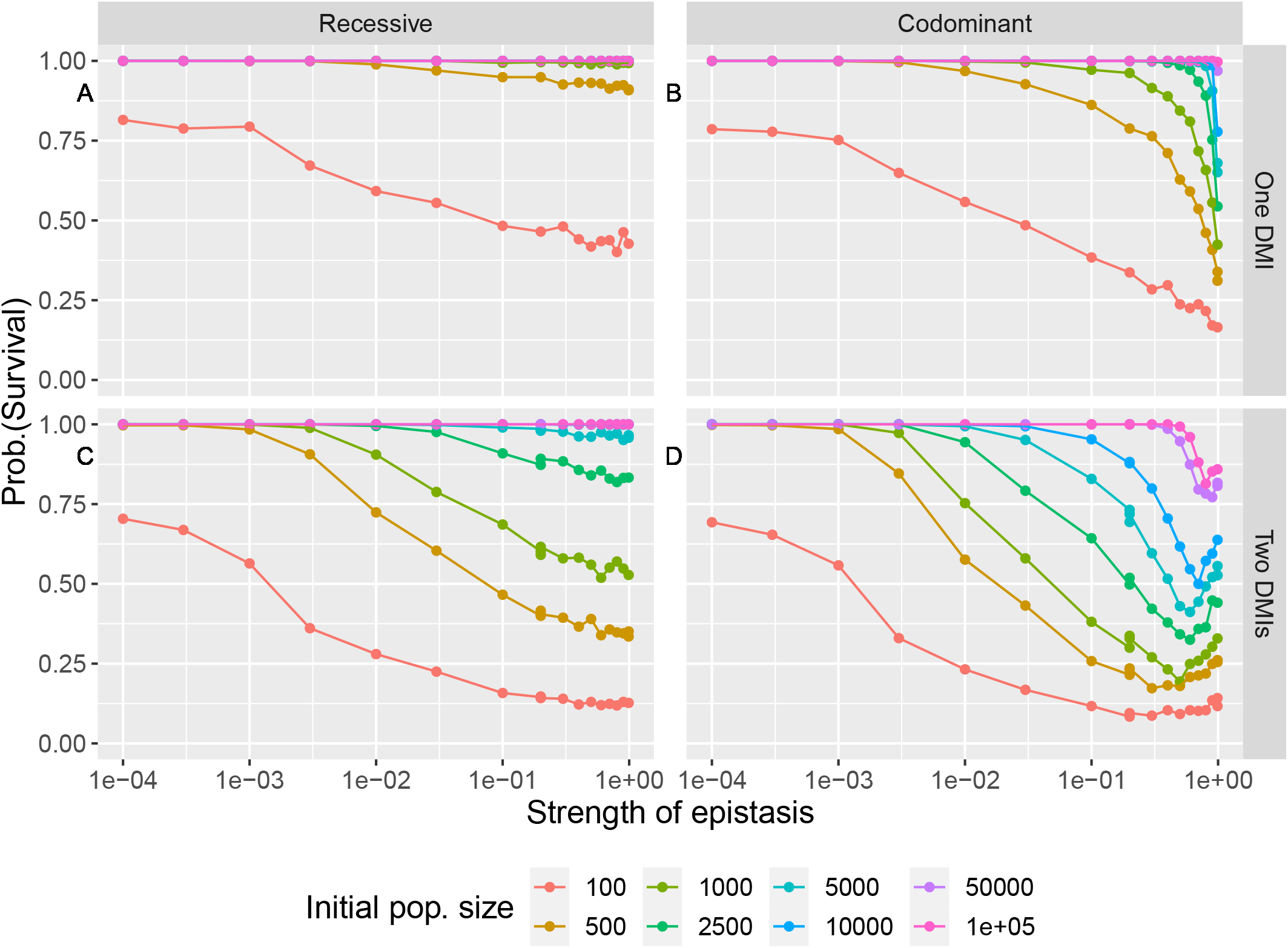
:The risk of extinction is not always a monotonous function of the model parameters. Here, all loci are freely recombining (*r* = 0.5). Lines serve as a guide for the eye. A) A single recessive DMI. B) A single codominant DMI. C) Two recessive DMIs. D) Two codominant DMIs. Survival probabilities are computed from 1,000 independent replicates. Error bars are not represented as their range is smaller or equal to the size of the data points. Other parameters are given in Table 1.

With two codominant DMIs, the population is at highest risk of extinction for intermediate values of epistasis (Fig. 2D). Here, the survival probability is non-monotonic because epistasis affects the hybrid population in two ways. On the one hand, epistasis initially reduces the population size due to the production of unfit individuals. More precisely, the growth rate of the population initially decreases below 1 as the strength of epistasis increases (see Fig. S2A for the F1 and F2 generations), which leads to a population size decline. This decline is less severe with increasingly weaker DMIs. On the other hand, the time to resolution of the genetic incompatibilities, at constant population size, decreases with the strength of epistasis (for *ϵ >* 0.001, Fig. S3). The resolution of the genetic incompatibilities relieves the population from the deleterious effects of the DMIs and the risk of extinction. Here, strong epistasis is favorable for survival. In summary, strong epistasis results in a short strong bottleneck, whereas weak epistasis results in a weak bottleneck of long duration. Populations with intermediate epistasis fall in between these two regimes, which leads to a longer period of time at a rather small population size (see Fig. S4). The longer time spent at low population sizes makes them more susceptible to extinction through drift. For recessive DMIs, the extinction probability strictly increases with the strength of the incompatibility (Fig. 2C). Since recessive DMIs are resolved much more slowly (Blanckaert and Bank, 2018), the regime of strong bottlenecks and fast resolution of the conflict does not exist here. The role of the size of the hybrid population at the time of the admixture event is independent of the dominance of the DMI: the smaller the population, the higher the risk of extinction (Fig. 2C-D).

**Table 1:** Parameters of the model and their default values. Parameters that varied in the main text are indicated with a *.

| Parameters | Notation | Default value |
| --- | --- | --- |
| Hybrid population size at the time of the admixture event | $N_{init}$ | 5,000 |
| Contribution of population A to the admixture event | $i_p$ | 0.5 |
| Selection coefficient for allele $a_k$ | $\alpha_k$ | -0.001 |
| Selection coefficient for allele $b_k$ | $\beta_k$ | -0.001 |
| Epistasis between allele $A_k$ and $B_k$ | $\epsilon_k$ | -0.2 |
| Mean number of offspring of the ancestral genotype | $\omega_{off}$ | 1.01 |
| Carrying capacity | $K$ | $10^6$ |
| Recombination rate between loci X and Y | $r_{XY}$ | 0.5 |
| Number of replicates | $n_{rep}$ | 1,000 |

In the absence of recombination or when it is rare, the linkage architecture does not affect the survival of the population (Fig. 3). Here, the initial population is at an unstable equilibrium at which only three genotypes exist: the two parental genotypes and the F1 hybrid genotype. Selection against F1s leads to a reduction in the population size until drift creates a sufficient frequency difference for selection to act against the rarer parental genotype. If F1 hybrids are (almost) non-viable, (Fig. S5), this regime of competition between the two parental genotypes always holds, regardless of the recombination rate and linkage architecture. This regime has three possible outcomes: the extinction of the hybrid population or the return to one of the two parental genotypes. The cost of producing F1 offspring drives the population towards extinction. Because epistasis creates positive frequency-dependent selection, departure from symmetry, generated by drift, ultimately leads to the resolution of the genetic incompatibilities, the fixation of one parental type (usually the more frequent), and the survival of the population. It is important to remark that the frequency dependence only applies to the marginal fitnesses of the alleles (and haplotypes); the fitness of the genotypes remains frequency-independent. If epistasis is recessive, the system is fully neutral in the absence of recombination. In this case, the presence of genetic incompatibilities, regardless of their strength, does not induce a risk of extinction. In the recessive scenario, a larger recombination rate increases the extinction probability, because recombination results in the formation of unfit genotypes. There is one notable exception (Fig S6 and S7, “Nested ABAB” architecture) where this effect is not monotonic, which we discuss below.

**Figure 3.**
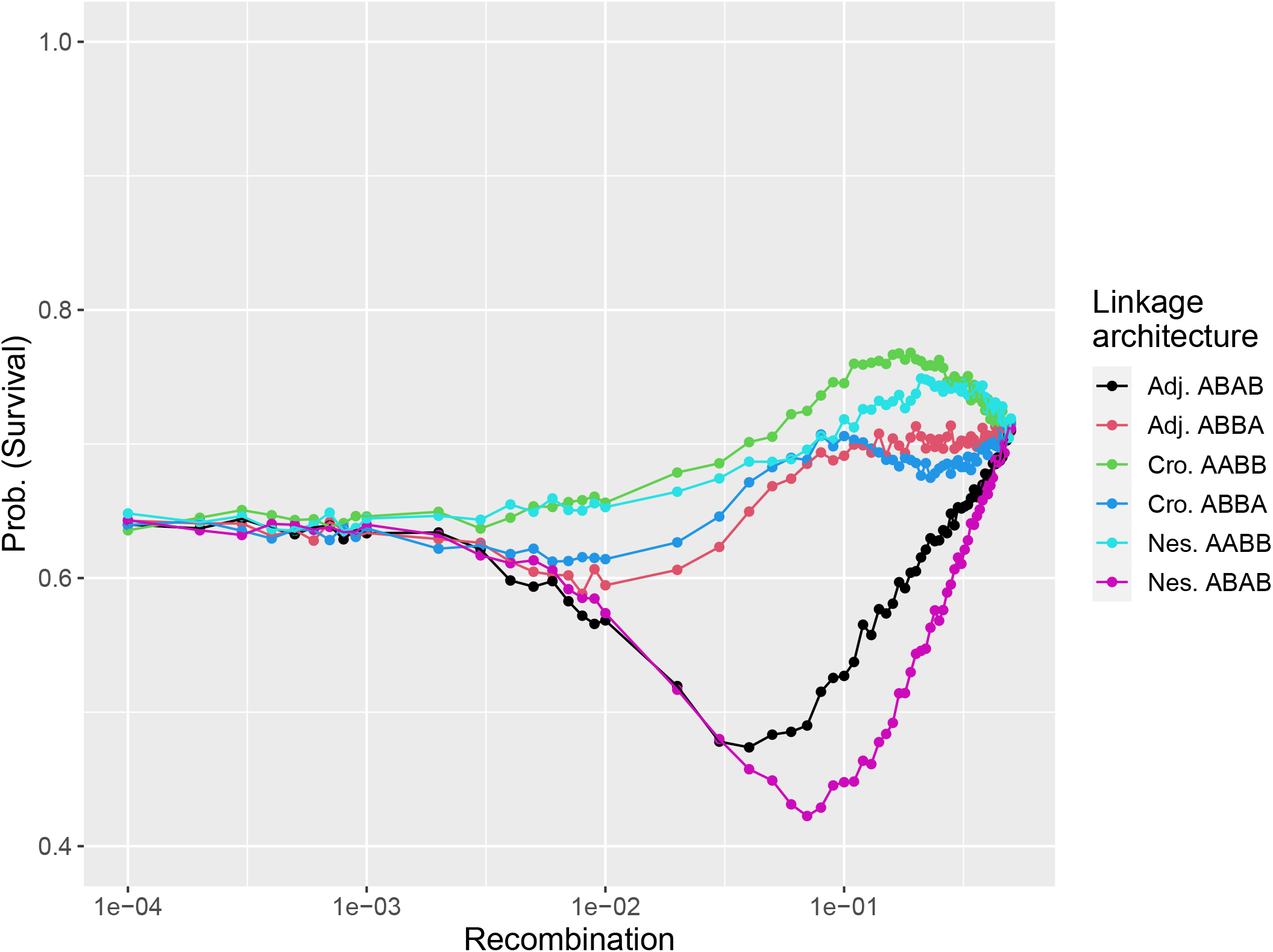
:The survival probability is often minimal for intermediate recombination rate and varies greatly between linkage architectures. We show the survival probability of the hybrid population as a function of recombination for the 6 different linkage architectures, for a pair of codominant DMIs of intermediate strength (*ϵ*_*k*_ = *−*0.2) and an initial population size *N*_init_ = 5, 000, estimated over 10,000 replicates. Lines serve as a guide for the eye and error bars are not represented as their range is smaller or equal to the size of the data points. The equivalent recessive case is shown in Fig. S5. Corresponding (both codominant and recessive) cases for lethal DMIs are shown in Fig. S6 and S7.

With codominant DMIs, the behavior is complex and non-monotonic; we here focus on the major patterns. A higher recombination rate can either increase the survival probability of the hybrid population (Fig. 3, “Crossed AABB” and “Crossed ABBA” architectures) or decrease it (“Adjacent ABAB” and “Nested AABB” architectures). Interestingly, Blanckaert and Bank (2018), in their soft selection model, observed a similar pattern for the probability of hybrid speciation, but for different sets of linkage architectures. To elucidate this architecture-dependent role of recombination, it helps to understand the dynamics in the absence of genetic drift. We first focus on the “Adjacent ABAB” and “Crossed AABB” architectures (Fig. 3, black and green dots). Both architectures generated similar strong probabilities of hybrid speciation for intermediate recombination rate in the soft selection regime (Blanckaert and Bank, 2018). In the present study, these two architectures behave very differently from each other (Fig. 3): the “Crossed AABB” architecture generates the highest chance of survival, and the “Adjacent ABAB” architecture the second lowest. One key difference between these two linkage architectures is the much higher frequency of the ancestral haplotype for the “Crossed AABB” architecture compared to the “Adjacent ABAB” architecture in the first generations of hybridization (Fig. S8). A higher frequency of the ancestral haplotype slows down the sorting of the DMIs, because parental haplotypes paired with the ancestral haplotype are safe from epistasis. When computing the marginal fitness of all 16 haplotypes (in the deterministic case, Fig. S9) for these two architectures, all but the ancestral haplotype show a higher marginal fitness for the “Crossed AABB” architecture case than in the “Adjacent ABAB” case. The initial higher frequency of the ancestral haplotype arises because the ancestral haplotype can be generated in the F2 breakdown with a single recombination event (Fig. 4 and S10). Conversely, it requires three recombination events to generate the ancestral haplotype (from the parental haplotypes) for the “Adjacent ABAB” architecture – a property that this linkage architecture shares with the “Nested ABAB” architecture, which displays the lowest survival rate. An excess of unfit genotypes, generated through the first recombination events, explains the low survival probability for the “Adjacent ABAB” and “Nested ABAB” architectures at intermediate recombination rates.

**Figure 4.**
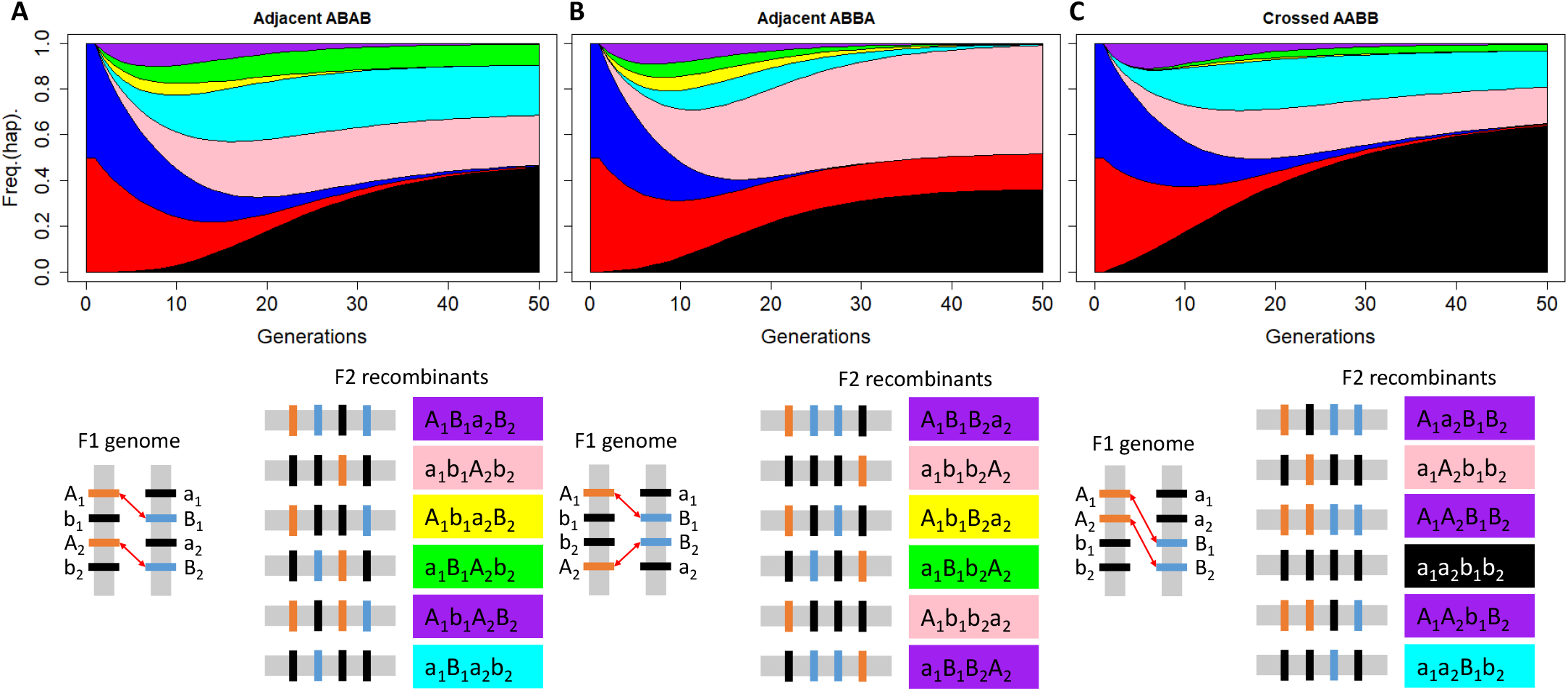
:Deterministic dynamics for the “Adjacent ABAB” (panel A), “Adjacent ABBA” (panel B), and “Crossed AABB” (panel C) architectures and the recombinant haplotypes formed in the F2 through single recombination event are indicative of the evolutionary fate of the population. The color indicates a haplotype or a group of haplotypes: the ancestral haplotype, *a*_1_*b*_1_*a*_2_*b*_2_, in black, the parental haplotypes in red (*A*_1_*b*_1_*A*_2_*b*_2_) and blue (*a*_1_*B*_1_*a*_2_*B*_2_), the ancestral haplotypes with a single derived allele in pink (*A*_1_*b*_1_*a*_2_*b*_2_ and *a*_1_*b*_1_*a*_2_*b*_2_) and cyan (*a*_1_*b*_1_*a*_2_*b*_2_ and *a*_1_*b*_1_*a*_2_*b*_2_), the hybrid haplotypes in yellow (*A*_1_*b*_1_*a*_2_*B*_2_) and green (*a*_1_*B*_1_*A*_2_*b*_2_), and the haplotypes harboring one or more incompatibilities in purple. On the right of the figure, we show the 6 possible recombinant haplotypes that can be formed through a single recombination event highlighted using the same color scheme. The deterministic dynamics were obtained for the same parameter values as Figure 3, with *r* = 0.1.

The F2 breakdown, which explains the difference between the two best and the two worst architectures in the codominant case, also sheds light on why only the “Nested ABAB” architecture displays a non-monotonic survival probability in the recessive case (Fig. S6 and S7). For all 6 linkage architectures, a single crossover forms two haplotypes that contain three ancestral alleles and one derived allele. For “Adjacent ABAB” and “Crossed AABB”, which have the highest survival probabilities for intermediate recombination rates, the emerging haplotypes, *a*_1_*A*_2_*b*_1_*b*_2_ and *a*_1_*a*_2_*B*_1_*b*_2_, are interaction-free because the two derived alleles stem from different DMIs. Conversely, for the two “Nested” architectures, the emerging haplotypes *a*_1_*A*_2_*b*_1_*b*_2_ and *a*_1_*a*_2_*b*_1_*B*_2_ contain an interaction between *A*_2_ and *B*_2_. This DMI generates a fitness cost for the hybrid population, which leads to a lower initial growth rate and, as a consequence, a lower survival probability. Interestingly, only the “Nested ABAB” architecture displays a minimum survival probability for intermediate recombination in the recessive case (Fig. S6 and S7). Here, the F3 recombining haplotype *a*_1_*A*_2_*b*_1_*B*_2_ is deleterious if homozygous, whereas its counterpart for the “Crossed AABB” architecture, *a*_1_*A*_2_*B*_1_*b*_2_ is always advantageous during the early generations of hybridization. Although this difference matters little for the codominant case, as the F1, as well as the F2 recombining haplotypes, are already expressing the incompatibilities, it becomes important in the recessive case, when the incompatibilities are not yet expressed. This difference leads to the distinct behaviors observed between the “Crossed AABB” and “Nested ABAB” architectures in the recessive case. Overall, the differences in survival probability highlight the influence of the linkage architecture on the effective strength of selection. Indeed, selection here is mainly epistatic and therefore frequency-dependent, and the linkage architecture controls which haplotypes initially form and at which frequencies.

Finally, in very small populations, recombination almost always reduces the probability of survival (in the codominant case; Fig. S11) and the difference between linkage architectures are far less pronounced. Drift erases most of the variance in model behavior that may arise from small differences in fitness, such as the small fitness advantages generated by forming “good” haplotypes in the F2 generation. In addition, the intrinsic disadvantage of the ancestral alleles over their derived counterpart barely changes the qualitative patterns (Fig. S12). The overall survival probability is increased when setting *α*_*i*_ and *β*_*i*_ to 0 (compared to *α*_*i*_ = *β*_*i*_ = *−*0.001), since the initial bottleneck is then less severe. Here, the effects of linkage architecture and recombination are slightly more pronounced than in the presence of direct selection. Both for the “Adjacent ABAB” and “Nested ABAB” architectures (and only these two), the pattern observed for larger populations is different from the others: the lowest survival probability is obtained for intermediate recombination rates in the codominant case.

### Resolution of the DMIs and survival probability

Both the strength of the bottleneck and its duration, caused by the resolution of the genetic incompatibilities, play a large role in the fate of the hybrid population. Figure 5 displays the relationship between the mean minimum population size (due to the bottleneck) and the survival probability. Here, the linkage architecture has little impact on the survival of the population. Although the linkage architecture determines the effective strength of selection (since it influences haplotype frequencies, and epistatic selection is frequency-dependent) and therefore the strength of the bottleneck, it does not impact the survival probability directly. In addition, for a given minimum population size, faster resolved incompatibilities are associated with an overall higher survival probability of the population, but the effect is marginal compared to the effect of the minimum population size itself (see also Fig. S13). The observed relationship between minimum population size and survival probability is similar between recessive and codominant incompatibilities (Fig. S14).

**Figure 5.**
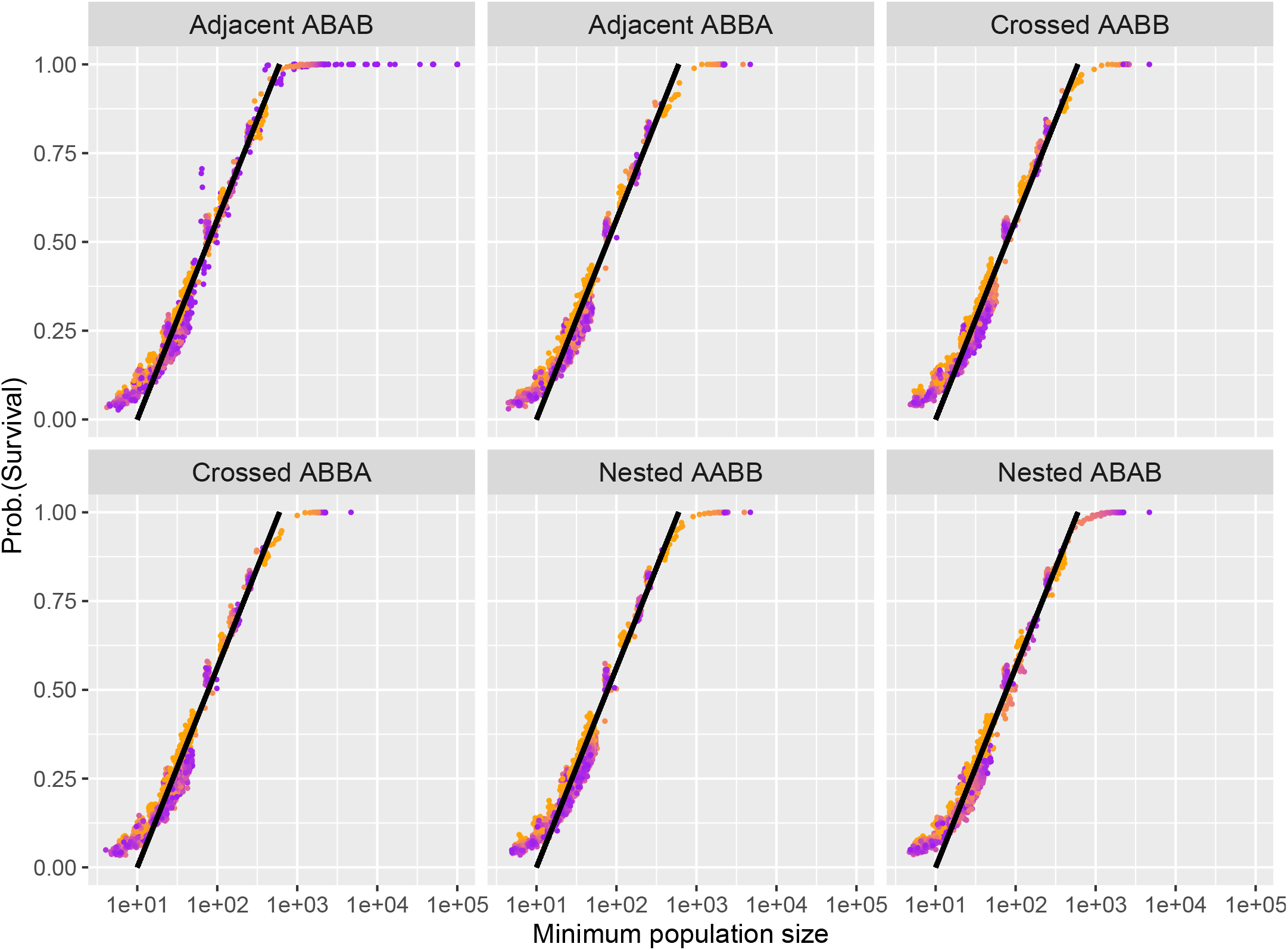
: The probability of survival is dictated by the minimum population size reached during the sorting of the (codominant) genetic incompatibilities. We show the survival probability for all 6 linkage architectures, for various initial population sizes, recombination rates and epistasis strength as a function of the minimum population size reached during the evolution. The black line is a visual guideline (y=-1/59+x/590) to facilitate comparison. Color indicates the recombination rate between loci (from *r* = 0 in orange to *r* = 0.5 in magenta; the equivalent recessive scenario is illustrated in Fig. S14.)

We found a similar relationship between minimum population size and survival probability in a classical evolutionary rescue scenario without incompatibilities (Fig. S15). More precisely, we considered a population doomed to extinction because of a negative growth rate, which could be rescued by a single beneficial mutation (Orr and Unckless, 2008). We set the growth rate 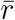 to that of the hybrid F1 population, and the selection coefficient *s* was chosen such that the growth rate of the final genotype matched that of the incompatibility-free population (Fig. S16). The survival probability for a given minimum population size is minimally higher in the classical evolutionary rescue scenario than in the hybrid population case (Fig S15). This is likely due to the difference in dynamics between the two scenarios: in the evolutionary rescue scenario, the mean growth rate increases over time, whereas it initially decreases during the sorting of the DMIs (as illustrated in Fig S2).

### Recombination (still) generates a Goldilocks zone for hybrid speciation

We next address the probability of hybrid speciation in the hard selection model. We recover qualitatively the main result observed for hybrid speciation with soft selection (Blanckaert and Bank, 2018): specific linkage architectures make hybrid speciation much more likely (Fig. 6). This holds independently of whether we do or do not condition on survival (Fig. 6). For intermediate recombination rates, and for 2 specific linkage architectures, hybrid speciation becomes a highly likely outcome. The underlying mechanism is identical to the soft selection regime. For intermediate recombination rates, these two architectures, “Adjacent ABAB” and “Crossed AABB” allow for allele *B*_1_ and *A*_2_ to escape into the ancestral background through a single crossover. The early formation of these haplotypes gives alleles, *B*_1_ and *A*_2_, a marginal fitness advantage over *A*_1_ and *B*_2_. These fitness differences, generated by the F2 breakdown, lead to a preferential reciprocal sorting of the DMIs towards *A*_2_ and *B*_1_ and therefore, per our definition, to hybrid speciation.

**Figure 6.**
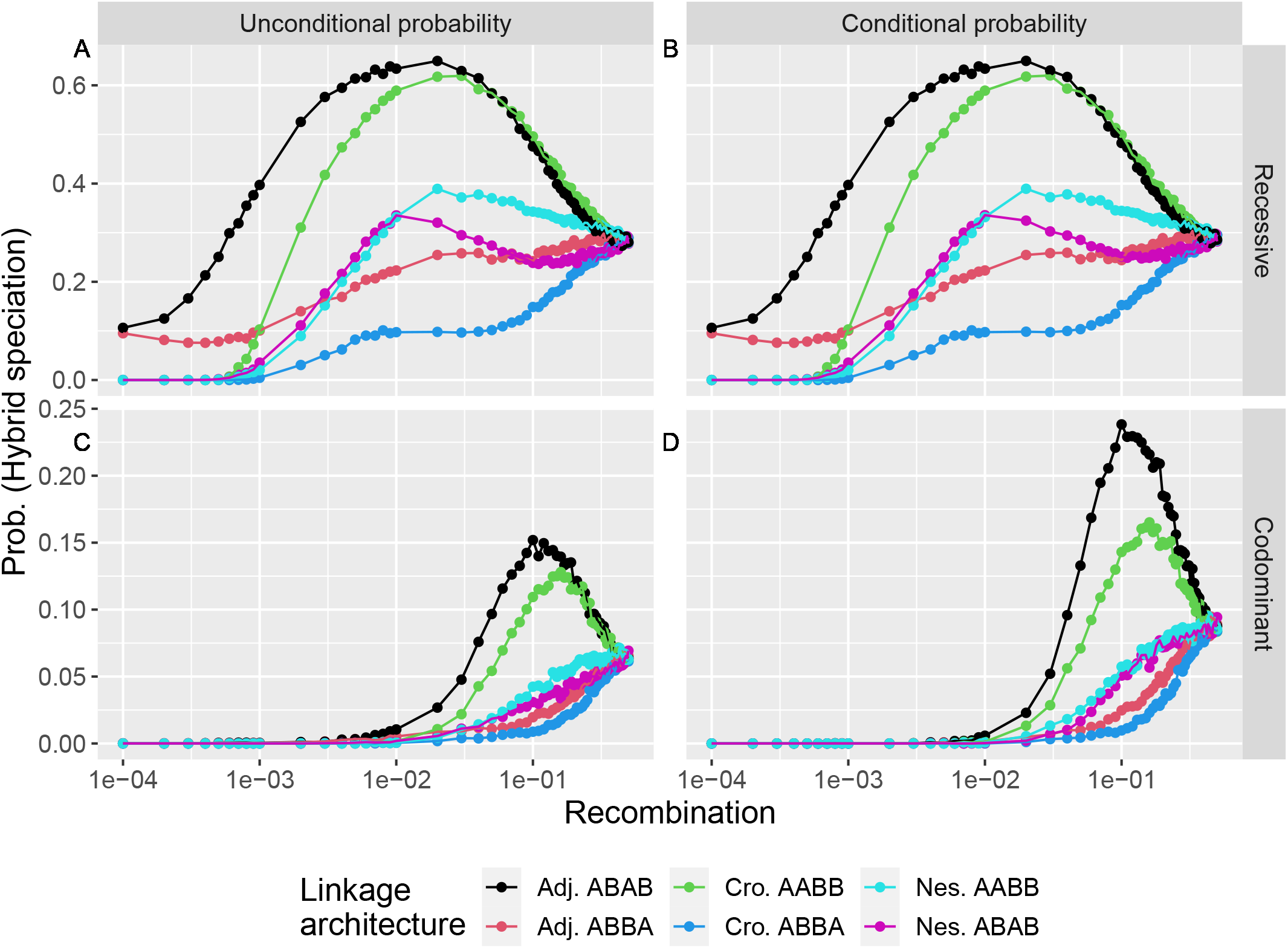
:The hybrid speciation probability is maximized at intermediate recombination rates for specific linkage architectures. We represent here the probability of hybrid speciation, as a function of the genetic distance between loci for all linkage architectures, with an initial population size *N*_init_ = 5, 000 measured over 10 000 replicates. Panel A) shows the hybrid speciation probability for recessive DMIs (*ϵ*_*k*_ = *−*0.2), panel B) shows the hybrid speciation probability, conditioned on survival, for recessive DMIs (*ϵ*_*k*_ = *−*0.2), panel C) shows the hybrid speciation probability for codominant DMIs (*ϵ*_*k*_ = *−*0.2) and panel D) shows the hybrid speciation probability, conditioned on survival, for codominant DMIs (*ϵ*_*k*_ = *−*0.2). Error bars are not represented as their range is smaller or equal to the size of the data points.

Although we have established that the minimum population size is a good predictor of the survival of the hybrid population, it is a very poor predictor of hybrid speciation. More precisely, it is an unreliable predictor since a given value for minimum population size corresponds to multiple hybrid speciation probabilities (Fig. S17). In particular, for the “Adjacent ABAB” and the “Nested ABAB” linkage architecture, both hybrid speciation (Fig. 6) and minimum population size (Fig. S18) are non monotonic functions of recombination: hybrid speciation is maximized for intermediate values of *r* and survival probability is minimized for similar intermediate values of *r*, creating a bifurcation pattern. This pattern was also observed for recessive DMIs (Fig. S19 and Fig. S20), indicating that the minimum population size is unrelated to the probability of hybrid speciation.

### Population size: beyond the effect of stochasticity

The hybrid speciation probability, given survival, is not always a monotonic function of the initial population size (Fig. 7, for codominant DMIs and Fig. S21 for recessive DMIs) for specific linkage architectures. First, we consider cases in which the probability of hybrid speciation, given survival, increases monotonically with the initial population size. To obtain reciprocal sorting of the two genetic incompatibilities, the initial parental haplotypes need to be broken down by recombination, which is unlikely to happen in very small populations. For these populations, the main two possible outcomes are the fixation of either parental genotype (Fig. S22). An increase in population size provides more opportunities and more time for the relevant haplotypes to appear and spread in the population. However, an increase in population size also reduces the effect of stochasticity and leads to the convergence of the hybrid population towards its deterministic final state. For the two linkage architectures, “Adjacent ABBA” and “Crossed ABBA”, which display this non-monomorphic behavior, the deterministic outcome is the fixation of the parental haplotype *A*_1_*A*_2_*b*_1_*b*_2_. Hybrid speciation is therefore extremely unlikely for large populations. Consequently, this dual effect of population size leads to a maximum hybrid speciation probability reached at intermediate initial population sizes. To obtain reciprocal sorting, the population size needs to be large enough for the hybrid speciation haplotypes to form and invade. Alternatively, resolution of the genetic incompatibilities can happen first through the spread of two one-derived-allele haplotypes (e.g. *A*_1_*a*_2_*b*_1_*b*_2_ and *a*_1_*a*_2_*b*_1_*B*_2_), that can form the hybrid speciation haplotype by recombination. Yet reciprocal sorting is not the deterministic outcome, thus drift needs to remain strong enough for hybrid speciation to occur.

**Figure 7:**
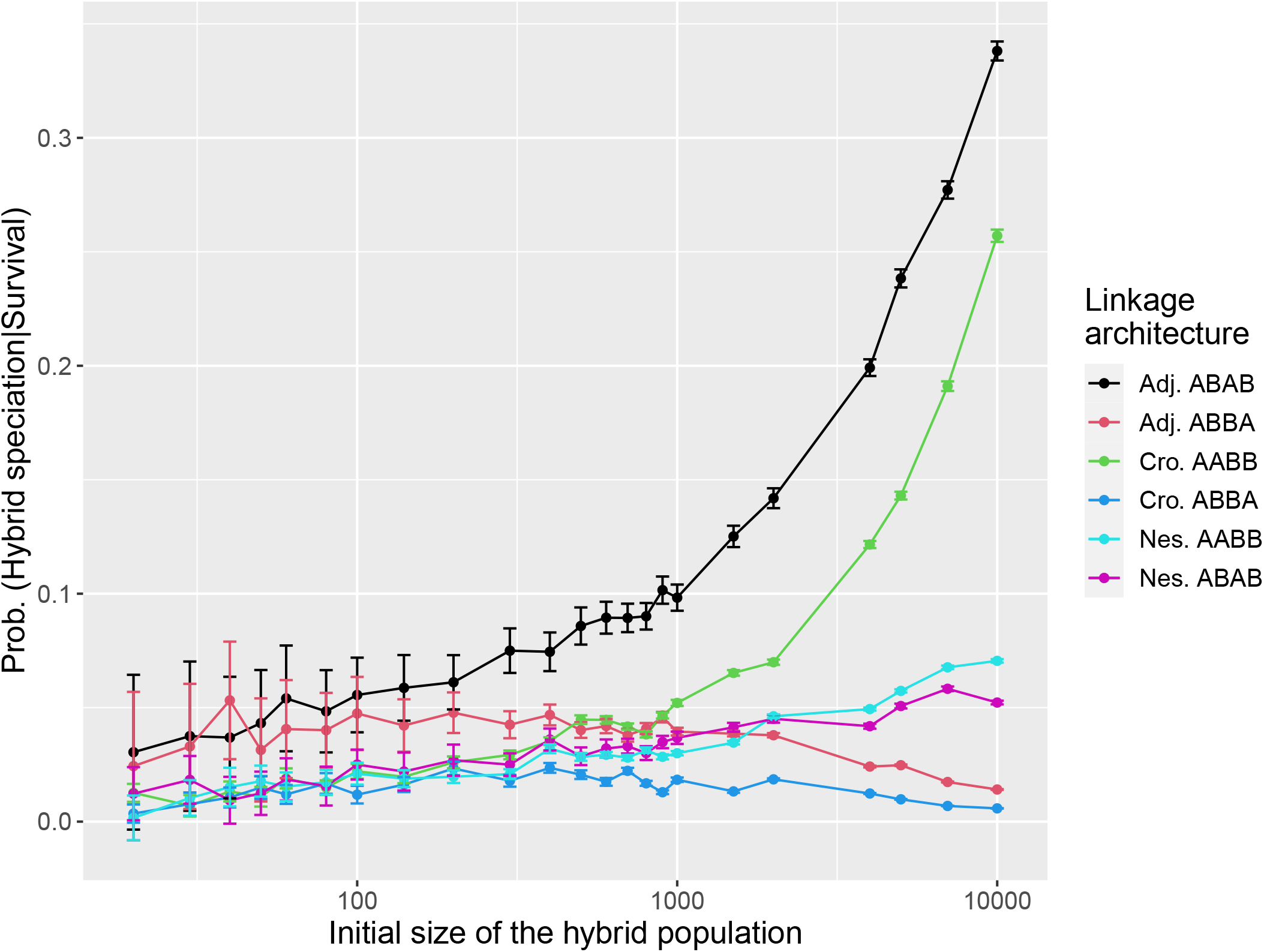
A larger initial population does not always facilitate hybrid speciation. We represent the probability of hybrid speciation, conditioned on survival, as a function of the initial population size for all linkage architecture for two pairs of codominant DMI (*ϵ* = *−*0.2) for intermediate recombination rates (*r* = 0.1) estimated from 10,000 replicates here. Lines serve as a guide for the eye and the errors bars correspond to the standard error. The underlying survival probabilities are illustrated in Figure S24.

The optimal linkage architecture to generate hybrid speciation changes depending on the influence of drift (Fig. 7 and Fig. S23). For small population sizes the two “Adjacent” linkage architectures display a higher probability of hybrid speciation given survival, whereas for large population “Adjacent ABAB” offers the best chance of observing hybrid speciation, whereas this outcome becomes impossible for “Adjacent ABBA”. We also observe this effect at low recombination rate for recessive DMIs (Fig. 7B).

Finally, the carrying capacity of the environment (*K*) neither alters the survival probability nor does it affect the hybrid speciation probability. We assumed that the carrying capacity of the environment was always quite large (*K* = 10^6^). As illustrated in the supplement (Fig S25-S27), reducing the carrying capacity of the environment to *K* = 5000 or even *K* = 500 does not alter the survival probability or the hybrid speciation probability given survival. This result supports our approach to neglect density dependence. Both the resolution of the hybrid incompatibilities and the survival of the population are largely determined during the initial phase following admixture, when population sizes are small. Therefore, by the time density regulation would set in, the evolutionary fate of the population has already been mostly decided. Furthermore, removing the intrinsic advantage of the derived allele over their ancestral counterpart (*α*_*k*_ = *β*_*k*_ = 0) does not change the hybrid speciation probabilities given survival (Fig. S28).

## Discussion

We investigated how the demographic and evolutionary dynamics of an admixed hybrid population impacted its fate. In particular, we were interested in two specific outcomes: the possibility of survival of the population and the reciprocal sorting of genetic incompatibilities, a mechanism suggested to initiate hybrid speciation. We focused on the role and effect of population size and linkage architecture on these two outcomes. We established that when multiple incompatibilities were involved, an intermediate strength of epistasis generated the highest risk of extinction for the newly formed population. Interestingly, the survival probability can be directly linked to the demographic dynamics of the population regardless of the linkage architecture or the parameters of the model. However, the demographic dynamics were affected by the underlying evolutionary parameters. The linkage architecture, i.e. the order of the incompatible loci along the genome, played a large role in determining the survival of the admixed population. It also had a considerable impact on the outcome of the reciprocal sorting of genetic incompatibilities and the possibility of hybrid speciation, as previously found in the soft selection scenario (Blanckaert and Bank, 2018). Yet, which linkage architectures were most favorable towards survival and reciprocal sorting was different. Linkage architectures favoring survival and hybrid speciation were different, suggesting that the probability of hybrid speciation is likely to be overestimated by models that ignored demographic dynamics. Finally, we highlighted the critical role of population size at changing the possible evolutionary outcomes: which linkage architecture was the most likely to generate hybrid speciation critically depended on the population size at the admixture event.

### Admixed populations and their parental sources

The genetic divergence between populations/species has long been suspected to play a large role in the fate of the hybrid population, and its possibility to evolve towards homoploid hybrid speciation (Abbott et al., 2013; Comeault and Matute, 2018) or recombinatorial speciation (Meier et al., 2019). The distribution of this divergence along the genome during the speciation process has also been largely debated, (islands of divergence Via (2012); Feder et al. (2012)) and whether reproductive isolation, in general, results from a few strongly differentiated regions or the resulting sum of many small effects (Flaxman et al., 2013, 2014). Blanckaert and Bank (2018) highlighted that reciprocal sorting of codominant DMIs may become the more difficult the more incompatibilities there are, due to the need of uncoupling the fate of different loci. Here, we add another consideration: the survival of the newly formed population during this sorting. The presence of multiple DMIs, both recessive and codominant, increases the risk of extinction of the population. This is true even if we control for the total amount of epistasis involved. The same F1 hybrid load, distributed across two freely recombining codominant DMIs instead of one, leads to lower survival (Fig. S29). In addition, we show that the extinction risk is maximized at intermediate strength of freely recombining genetic incompatibilities (Figure 2). This result has further implications on which populations may give rise to hybrid species. In the early stages of the speciation process (Feder et al., 2012), the parental populations may be partially isolated because of few islands of divergence harbouringweak unlinked hybrid incompatibilities. The further accumulation of weak DMIs in tight linkage to these islands (i.e. the gradual build-up of an island of divergence) is then expected to effectively increase the overall epistasis generated by the island of divergence as a whole and thereby the extinction risk of a new isolated hybrid population. By taking these two effects (i.e., splitting the hybrid load between multiple unlinked DMIs, and slowly increasing the strength of epistasis at a given region, as the parental populations diverge) into account, our result supports the idea that having multiple unlinked islands of divergence is unlikely to favor hybrid speciation (Blanckaert and Bank, 2018). The addition of the demographic dynamics to the evolutionary dynamics in our model strengthens the view that only recently diverged parental species may give rise to hybrid species following admixture. This result is different from the one obtained by Comeault and Matute (2018), but it is important to consider that mechanisms of reproductive isolation considered are different, assortative mating versus intrinsic incompatibilities, leading to this possible discrepancy. It was recently argued by Servedio and Hermisson (2020) that partial reproductive isolation may not only be a transient state, as it often assumed, but a possible evolutionary outcome itself. In such a scenario, partial reproductive isolation may help maintain the parental populations in a genetic and phenotypic state of “distant enough but not too distant”, making both hybrid speciation or even recombinatorial speciation (“a special form of hybrid speciation involving karyotype evolution,” Marques et al. (2019)) more likely than expected.

Finally, hybrid speciation depends on the nature of the event that forms the initial admixed population, and how the admixed population is connected to its parental sources. A classical hybrid zone with constant gene flow seems an unlikely candidate for hybrid speciation, as it would prevent the reciprocal sorting of the DMIs. Here, we have investigated a best case scenario: we assume a single colonization event through an admixed population, followed by isolation from both parental sources. This scenario can be caused by extraordinary climatic events that lead to rare long range dispersal of many individuals (e.g., a flood for fish, a storm for seed dispersal). Depending on the organisms involved, such events could involve a few individuals or 1000s. The founding population size directly influences the likelihood of hybrid speciation,as illustrated in Figure 7.

### Role of linkage architecture

The inclusion of the demographic dynamics of the hybrid population in our study further highlighted the importance of the linkage architecture, i.e. the order of loci along the chromosome. Previously, we found that 2 out of 6 linkage architectures were extremely favorable to reciprocal sorting at intermediate recombination distance between loci (Blanckaert and Bank, 2018).This result remains valid under a hard selection regime, but one of the two linkage architectures also resulted in one of the lowest survival probabilities. This phenomenon happens at the same genetic distance between loci that maximizes the chance of hybrid speciation. These patterns explain why hybrid speciation is repeatable in some lineages and impossible in others, and makes hybrid speciation an odd outcome: a “high risk, high reward” scenario. Overall, after including the demographic dynamics into our model, we suggest that hybrid speciation will happen for organisms that can generate rare but large bursts of migration, such that admixed populations quickly colonize new environments -yet without permanent gene flow. As such the formation of the Italian sparrow species (Runemark et al., 2018) could fit into this category. Also, the iconic adaptive radiation of ciclid fish observed in Lake Mweru was suspected to emerge from specific hybridization events, although further crosses between other sister species yield no hybrid species Meier et al. (2019). Despite many additional factors at play (e.g., local adaptation, niche availability), such observations are in agreement with our results.

### Are recessive DMI the key to hybrid speciation?

For recessive DMIs (i.e. no fitness cost for F1 hybrids), the presence of DMIs reduces the chance of survival of the population. However, this effect is generally much weaker than for codominant DMIs, and monotonic with the strength of the genetic incompatibilities. The monotonic relationship between survival and epistasis indicates that intermediate-strength barriers can contribute to hybrid speciation as suggested by Comeault and Matute (2018). Furthermore, the linkage architecture most likely to generate reciprocal sorting and hybrid speciation in the minimal model presented here (“Adjacent ABAB”) does not suffer from excess mortality (as opposed to the codominant case, where the “Adjacent ABAB” has a lower survival probability than most others linkage architectures). Overall, this suggests that any successful reciprocal sorting event is more likely to involve recessive DMIs than codominant DMIs. Given that there is both theoretical and empirical evidence, that most DMIs should be recessive (Orr, 1993; Presgraves, 2003), reciprocal sorting of multiple recessive DMIs may be the path to homoploid hybrid speciation.

Yet, recessive incompatibilities form poor barriers to gene flow, as shown for the soft selection regime (Blanckaert and Bank, 2018). In the model investigated here, a population will either go extinct or reach the carrying capacity of the environment. Unless this carrying capacity is really low, all final hybrid populations are equivalent to those in a soft selection scenario. Therefore, measures of reproductive isolation should be qualitatively equivalent to those obtained in Blanckaert and Bank (2018). The effective migration rate is only reduced for almost lethal epistasis and when the different loci are located far apart from each other. Overall, including the effect of hard selection reinforces the dichotomy already observed with soft selection. Reciprocal sorting of codominant DMIs, while quite unlikely, can effectively isolate a hybrid population from its parental sources. On the contrary, reciprocal sorting of recessive DMIs is far more likely but creates poor barriers to gene flow.

## Conclusion

The role of hybridization as source of potential adaptation has recently been highlighted (Marques et al., 2019). We demonstrated here the importance of considering both the demographic and evolutionary dynamics to study the fate of these admixed populations, since their actions can be antagonistic and the evolutionary outcomes strongly dependent on the population size. Overall, our results further confirm the importance of the linkage architecture, i.e. the order of the different loci along the genome, for hybrid speciation and also for the survival probability.

## Acknowledgments

We thank the Bank lab for discussion of the project. This study was funded by an ERC Starting Grant 804569 (FIT2GO) and HFSP Young Investigator Grant RGY0081/2020 to CB . The research was partly supported by National Institutes of Health (NIH) grant R35 GM139412 to Bret Payseur.

