## SupplementaryInformation for "In search of the Goldilocks zone for hybrid speciation II: hard times for hybrid speciation?"

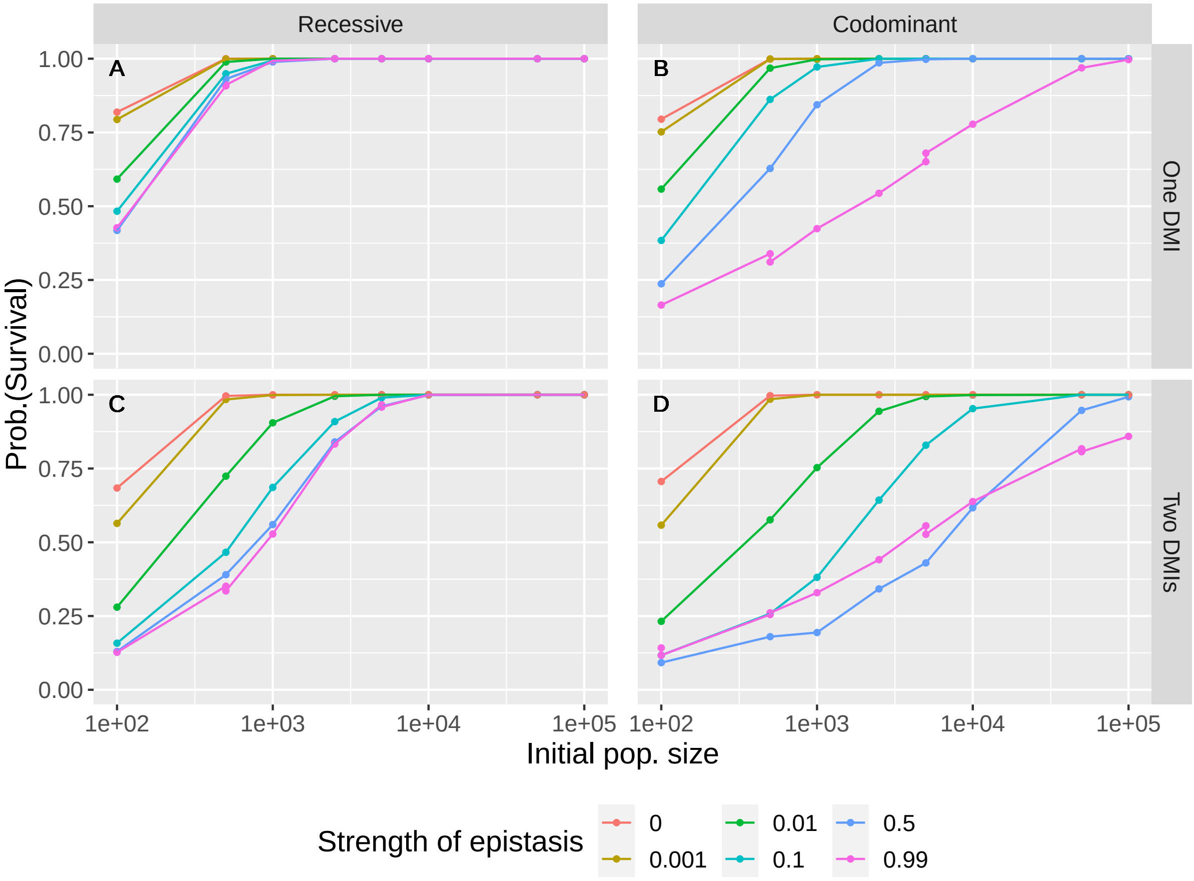


Figure S1: The survival probability of the hybrid population increases with the initial population size but intermediate strength of epistasis presents the highest risk of extinction. Here all loci are freely recombining. The lines correspond to local regressions and mainly serve to provide a guide for the eye. A) A single recessive DMI. B) A single codominant DMI. C) Two recessive DMI. D) Two codominant DMIs. Additional parameters are set to $n_{\text{rep}} = 1,000, N_{\text{init}} = 5,000, r = 0.5, \alpha_1 = \alpha_2 = \beta_1 = \beta_2 = -0.001, K = 10^6$.

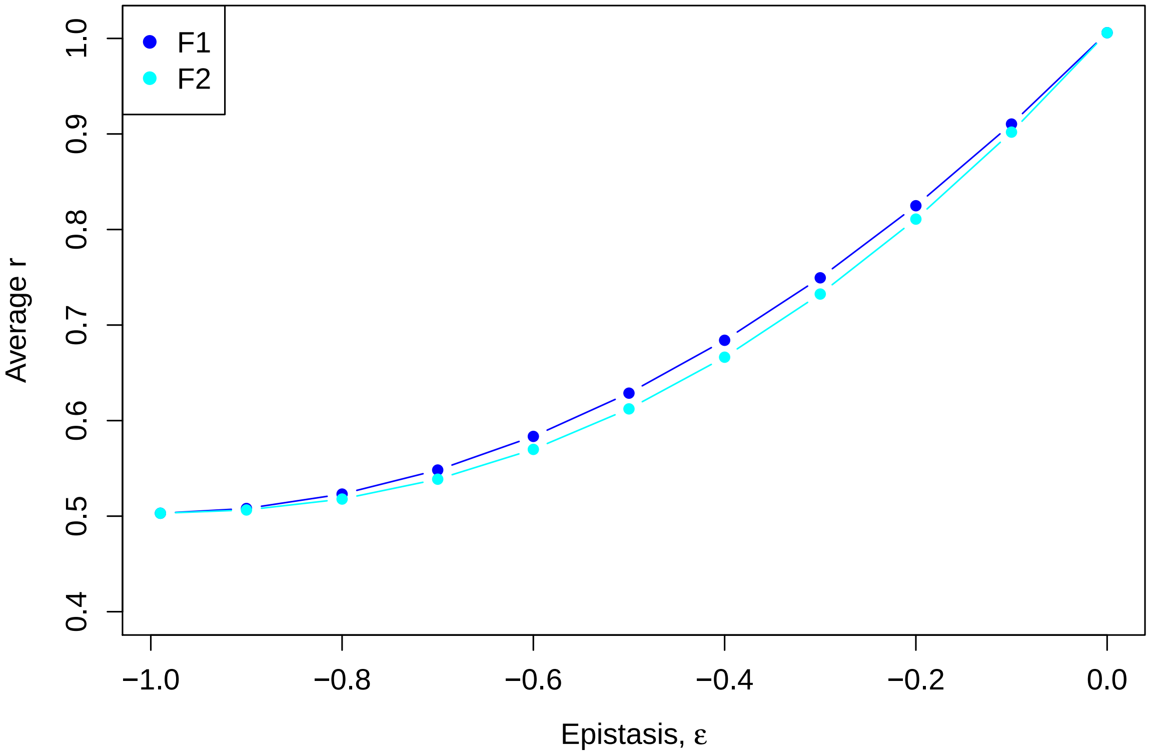


Figure S2: Average growth rate, $\bar{r}$ of the F1 and F2 generation for the hybrid population in the deterministic case, for two pairs of codominant DMIs and freely recombining loci. Additional parameters are set to $r = 0.5, \alpha_1 = \alpha_2 = \beta_1 = \beta_2 = -0.001$.

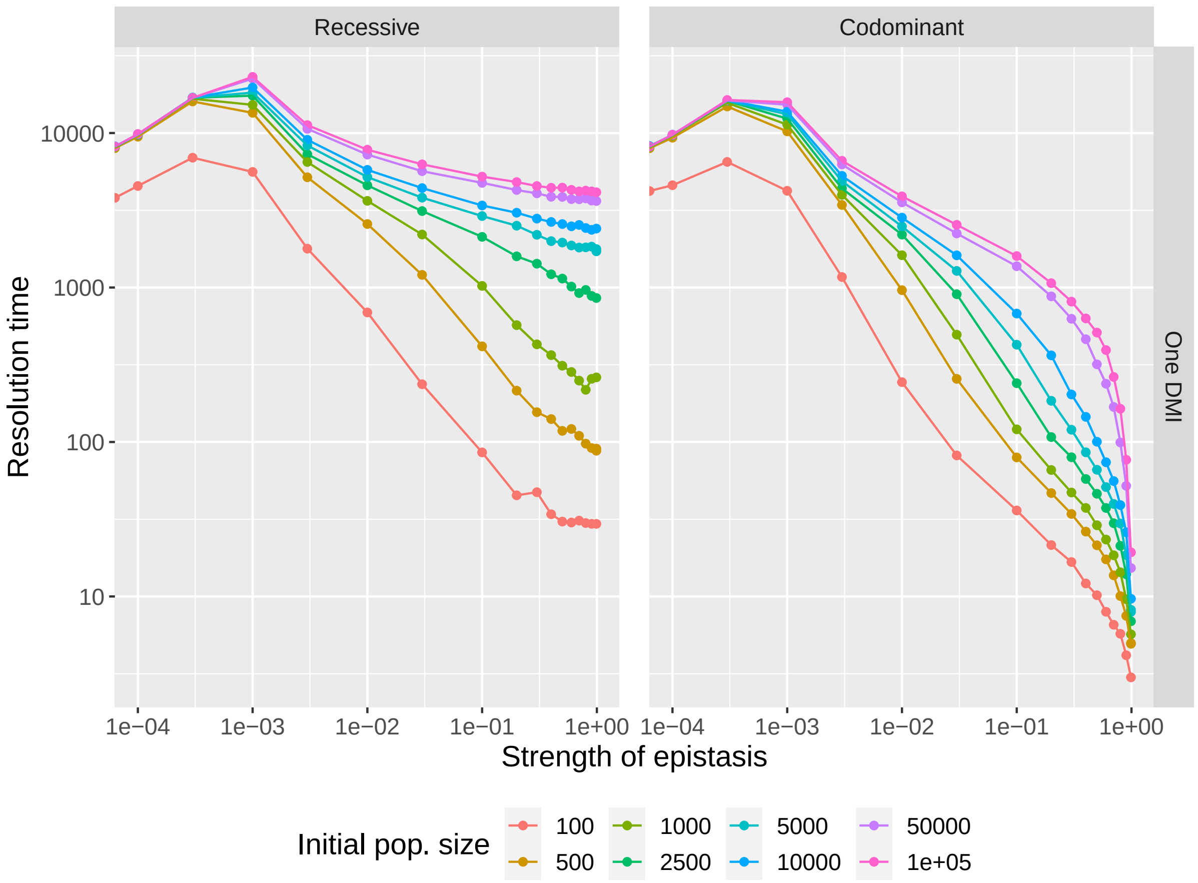


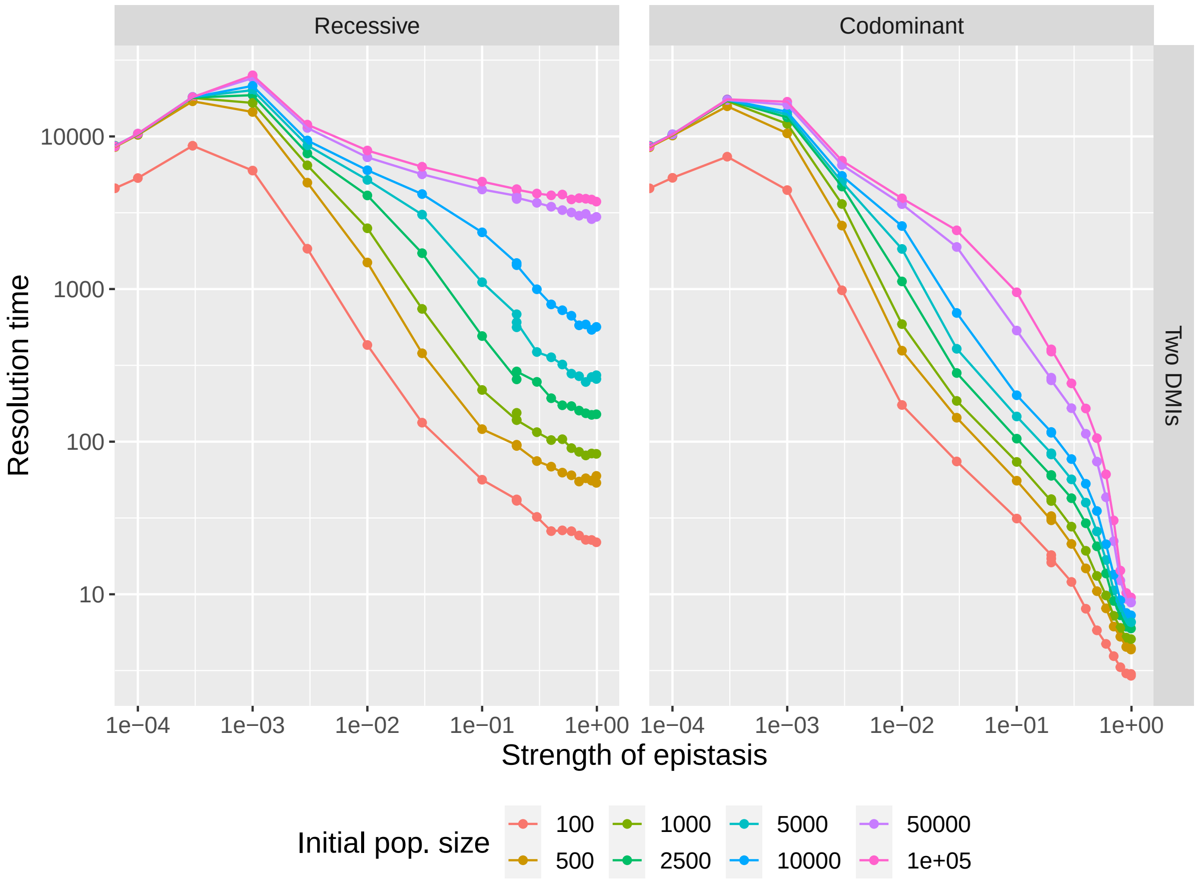


Figure S3: Mean resolution time of the genetic incompatibilities for recessive and codominant unlinked genetic incompatibilities. Additional parameters are set to $\alpha_1 = \alpha_2 = \beta_1 = \beta_2 = -0.001, n_{\text{rep}} = 1,000$

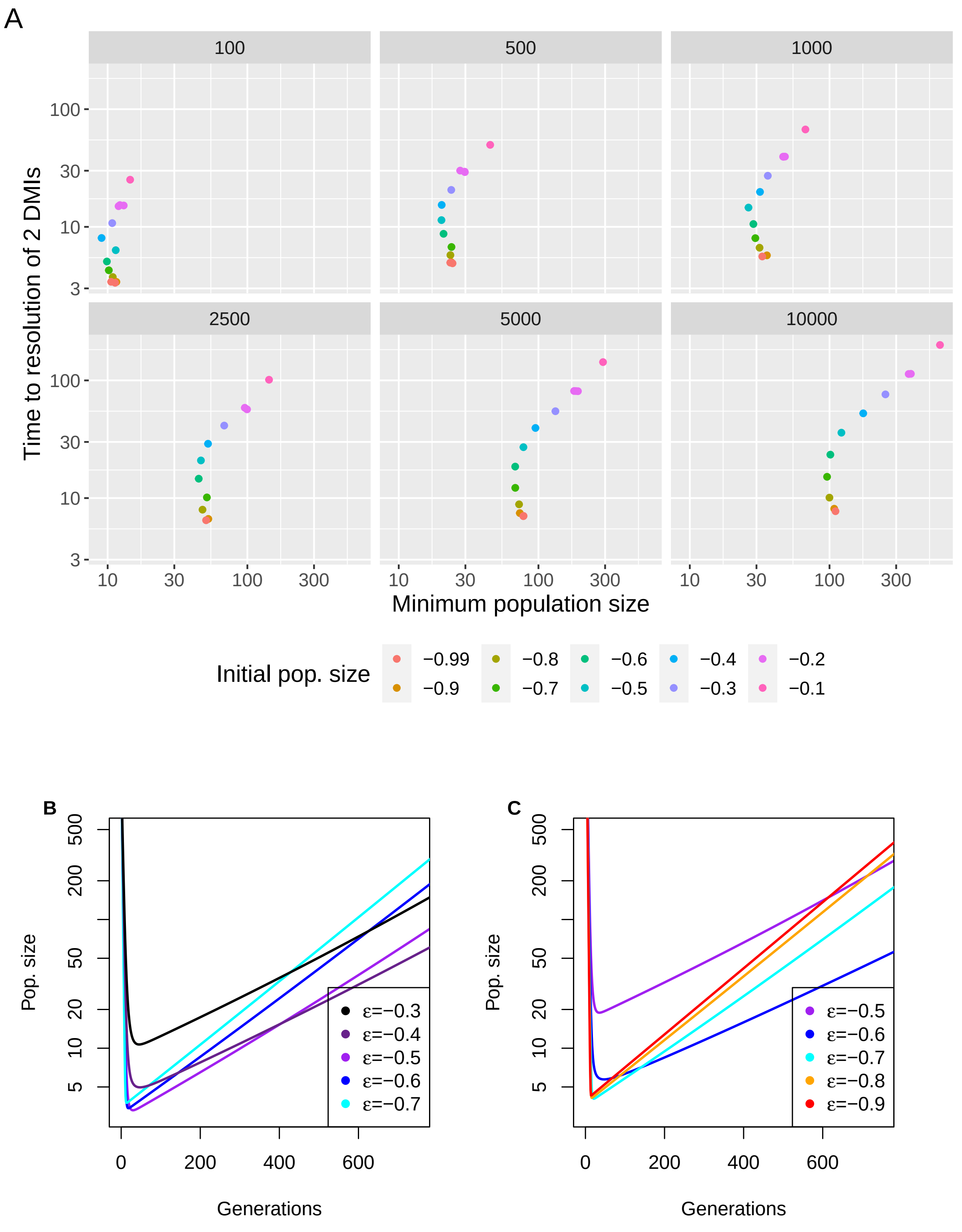


Figure S4: A) Time to resolve the 2 DMIs as a function of the minimum population size reached. Each color corresponds to a different epistasis coefficient and each panel to a different initial population size. B) Deterministic trajectories for the case $N_{\text{init}} = 500$ C) Deterministic trajectories for the case $N_{\text{init}} = 5000$. Additional parameters are set to $r = 0.5, \alpha_1 = \alpha_2 = \beta_1 = \beta_2 = -0.001, K = 10^6$.

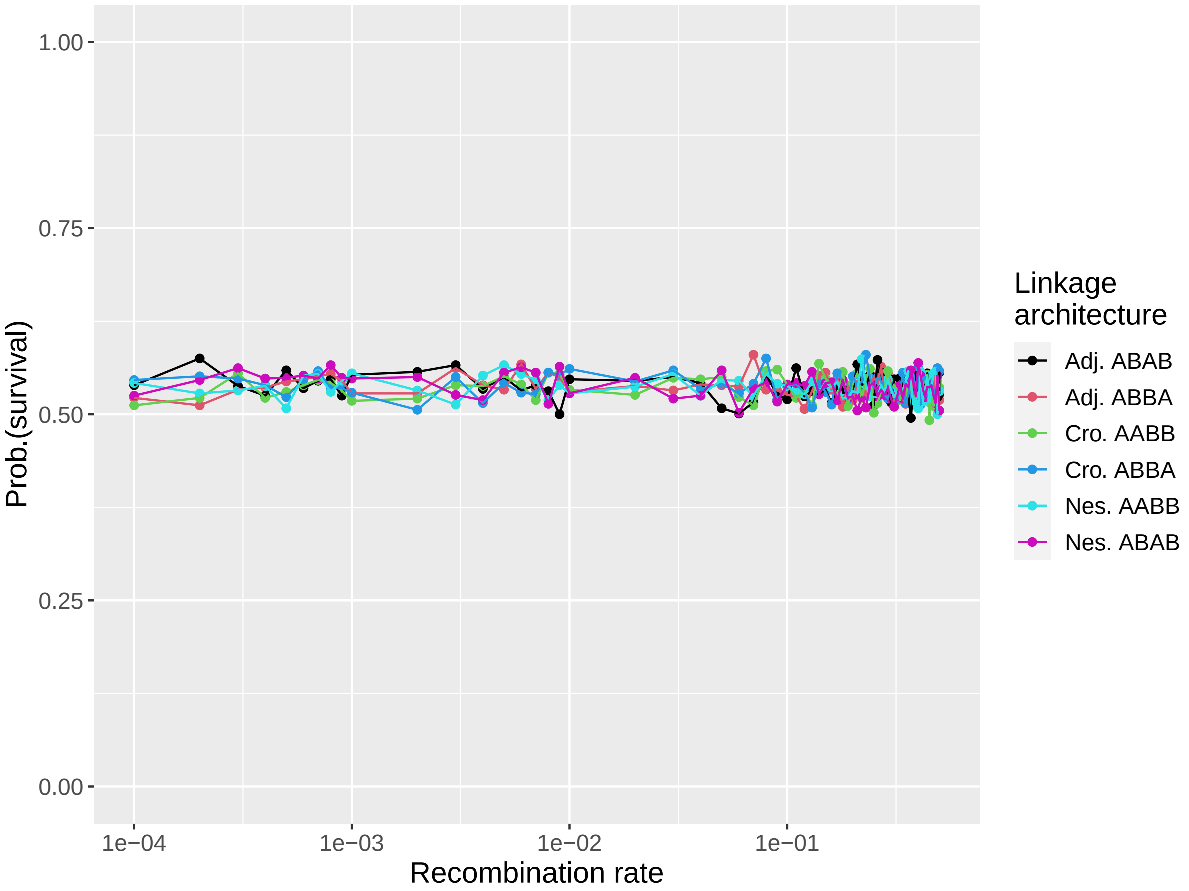


Figure S5: Survival probability as a function of recombination for all 6 architectures for the codominant (quasi-)lethal case ($\epsilon = -0.99$). Additional parameters are set to $N_{\text{init}} = 5,000, \alpha_1 = \alpha_2 = \beta_1 = \beta_2 = -0.001, K = 10^6, n_{\text{rep}} = 1,000$

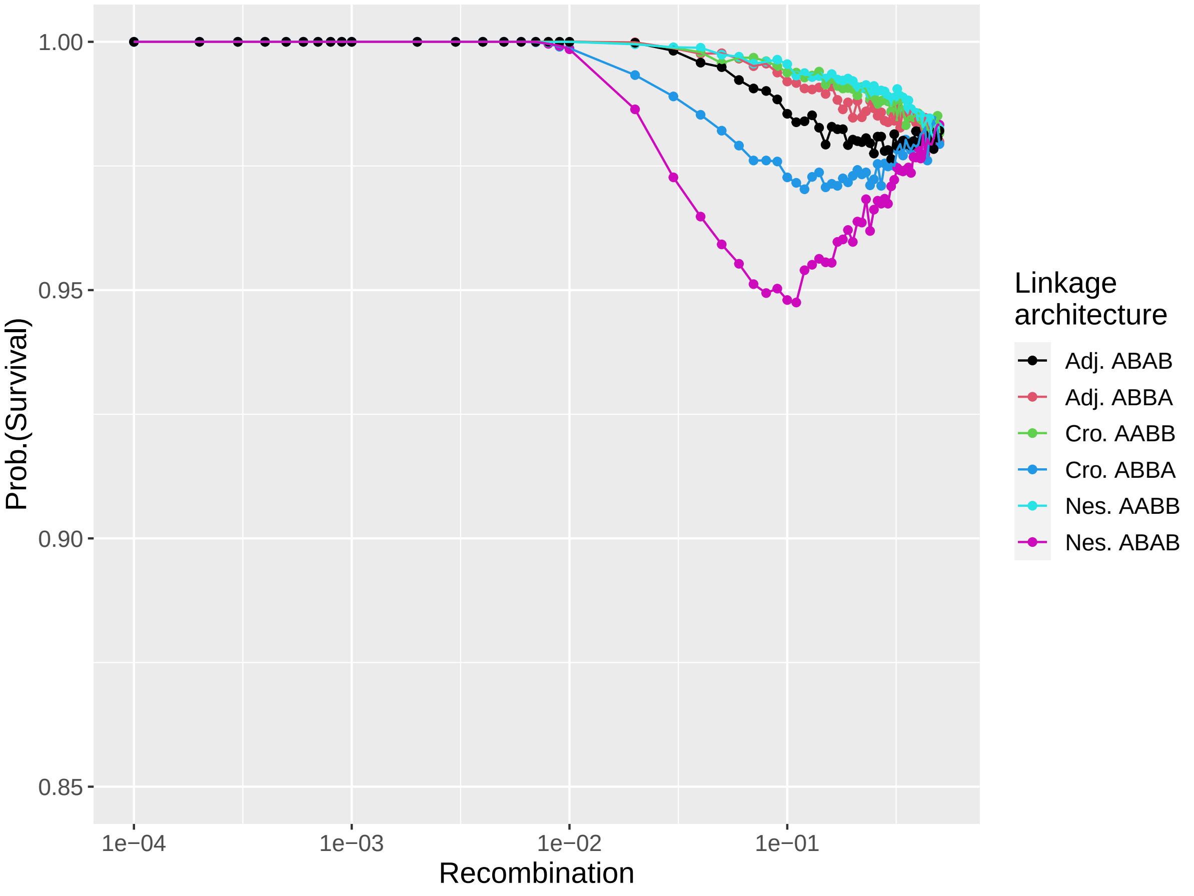


Figure S6: Survival probability as a function of recombination for all 6 architectures for the recessive case ($\epsilon = -0.2$). Additional parameters are set to $N_{\text{init}} = 5,000, \alpha_1 = \alpha_2 = \beta_1 = \beta_2 = -0.001, K = 10^6, n_{\text{rep}} = 10,000$

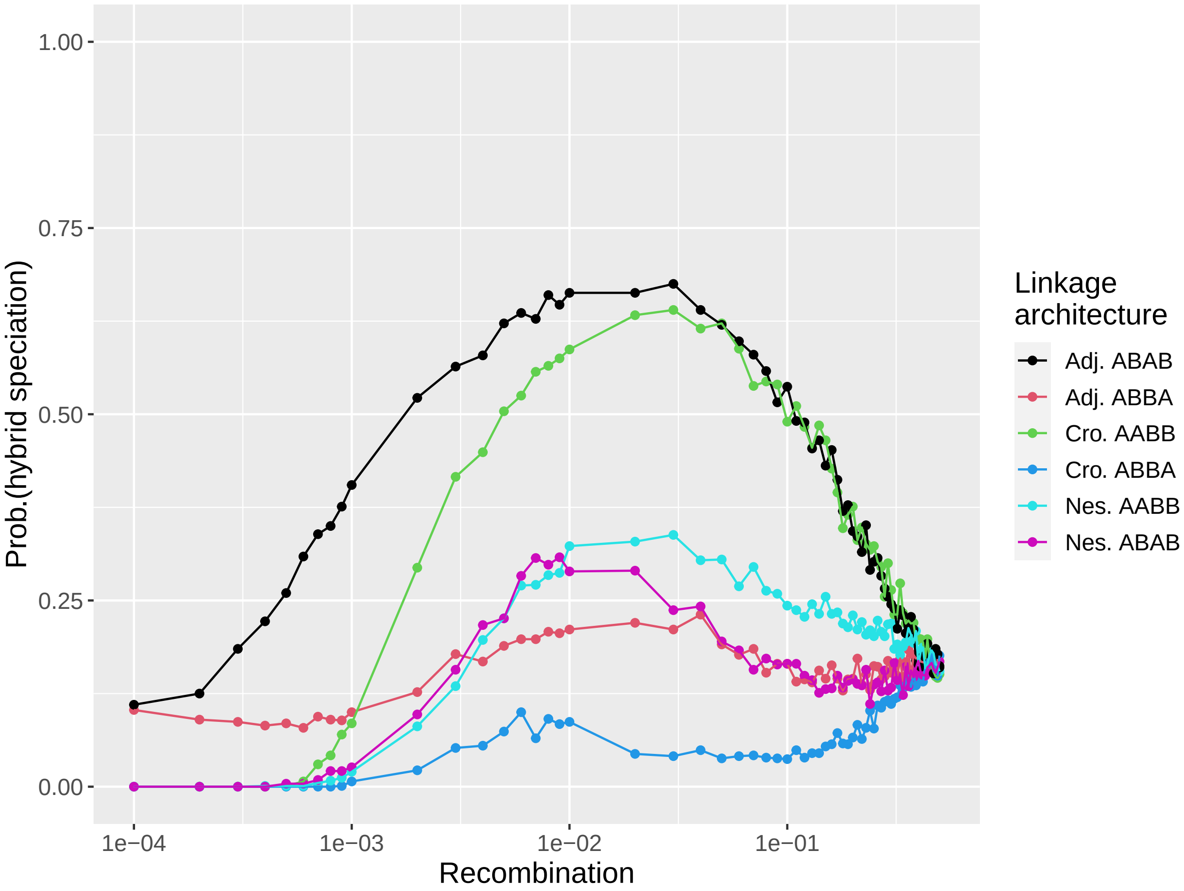


Figure S7: Survival probability as a function of recombination for all 6 architectures for the recessive (quasi-)lethal case ($\epsilon = -0.99$). Additional parameters are set to $N_{\text{init}} = 5,000, \alpha_1 = \alpha_2 = \beta_1 = \beta_2 = -0.001, K = 10^6, n_{\text{rep}} = 1,000$

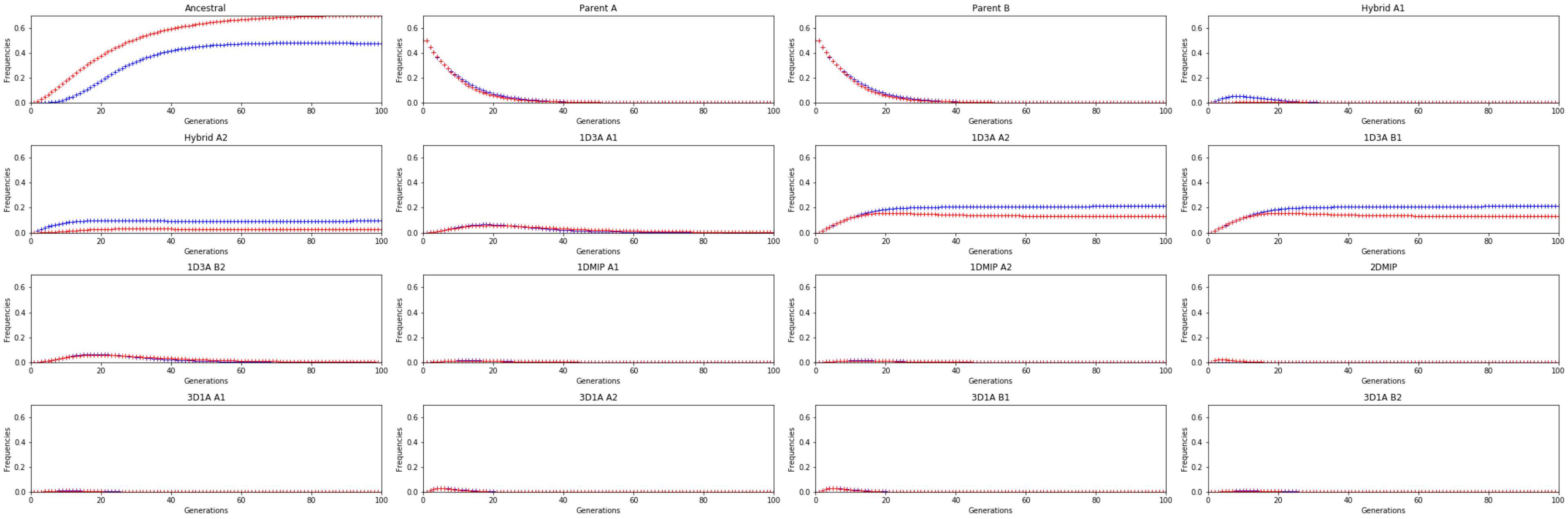


Figure S8: Frequency dynamics of all 16 haplotypes for the adjacent ABAB (blue) and crossed AABB (red) architectures in the deterministic case for $r = 0.1$. Additional parameters are set to $\alpha_1 = \alpha_2 = \beta_1 = \beta_2 = -0.001, K = 10^6$

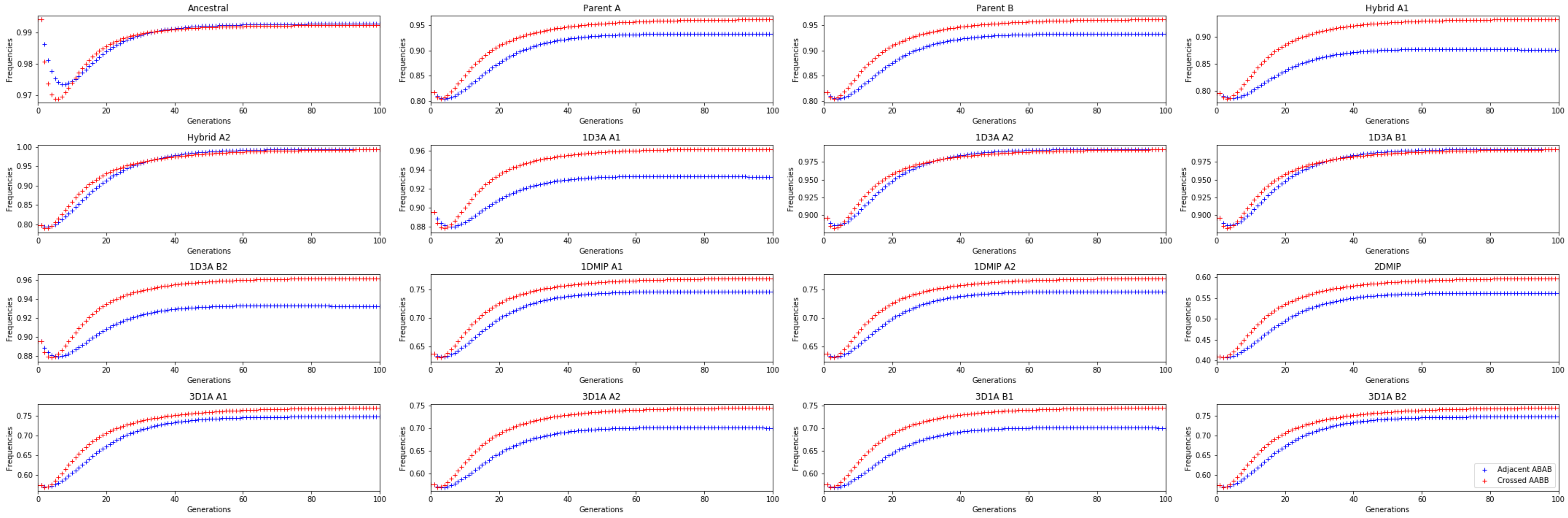


Figure S9: Marginal fitness of all 16 haplotypes for the adjacent ABAB (blue) and crossed AABB (red) architectures in the deterministic case for $r = 0.1$. Additional parameters are set to $\alpha_1 = \alpha_2 = \beta_1 = \beta_2 = -0.001, K = 10^6$

| **Linkage architecture** | **F1 hybrid** | **F2 recombinant resulting from a single recombining event between the** | | | **Remarkable properties** |
|---|---|---|---|---|---|
| | | 1st and 2nd loci | 2nd and 3rd loci | 3rd and 4th loci | |
| Adjacent ABAB | 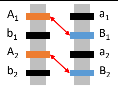 | $A_1B_1a_2B_2$<br>$a_1b_1A_2b_2$ | $A_1b_1a_2B_2$<br>$a_1B_1A_2b_2$ | $A_1b_1A_2B_2$<br>$a_1B_1a_2b_2$ | Bias towards B1 and A2 |
| Adjacent ABBA | 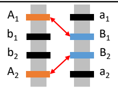 | $A_1B_1B_2a_2$<br>$a_1b_1b_2A_2$ | $A_1b_1B_2a_2$<br>$a_1B_1b_2A_2$ | $A_1b_1b_2a_2$<br>$a_1B_1B_2A_2$ | Bias towards A1 and A2 |
| Crossed AABB | 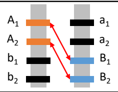 | $A_1a_2B_1B_2$<br>$a_1A_2b_1b_2$ | $A_1A_2B_1B_2$<br>$a_1a_2b_1b_2$ | $A_1A_2b_1B_2$<br>$a_1a_2B_1b_2$ | Bias towards B1 and A2<br>Ancestral hap. in 1 step |
| Crossed ABBA | 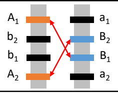 | $A_1B_2B_1a_2$<br>$a_1b_2b_1A_2$ | $A_1b_2B_1a_2$<br>$a_1B_2b_1A_2$ | $A_1b_2b_1a_2$<br>$a_1B_2B_1A_2$ | Bias towards A1 and A2 |
| Nested ABAB | 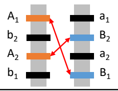 | $A_1B_2a_2B_1$<br>$a_1b_2A_2b_1$ | $A_1b_2a_2B_1$<br>$a_1B_2A_2b_1$ | $A_1b_2A_2B_1$<br>$a_1B_2a_2b_1$ | |
| Nested AABB | 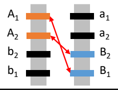 | $A_1a_2B_2B_1$<br>$a_1A_2b_2b_1$ | $A_1A_2B_2B_1$<br>$a_1a_1b_2b_1$ | $A_1A_2b_2B_1$<br>$a_1a_2B_2b_1$ | Ancestral hap. in 1 step |

Figure S10: Recombining gametes produced by F1 individuals assuming a single recombination event per genome. In the last column, we denote remarkable features of specific architectures, that are linked to evolutionary outcomes: a bias in the sorting of the DMIs towards one of the two following allele pairs $A_1, B_2$ or $A_2, B_1$ leads to a higher chance of hybrid speciation; a bias towards allele of the same origin (e.g., $A_1, A_2$) leads to a resolution towards the corresponding parental species, and as discussed in Blanckaert and Bank (2018), lead to hybrid speciation being maximized for asymmetric contribution of the parental species. Finally, the ability to quickly generates the anestrla haplotype acts as a buffer against extinction risk. This figure is adpated from Figure 6 in Blanckaert and Bank (2018).

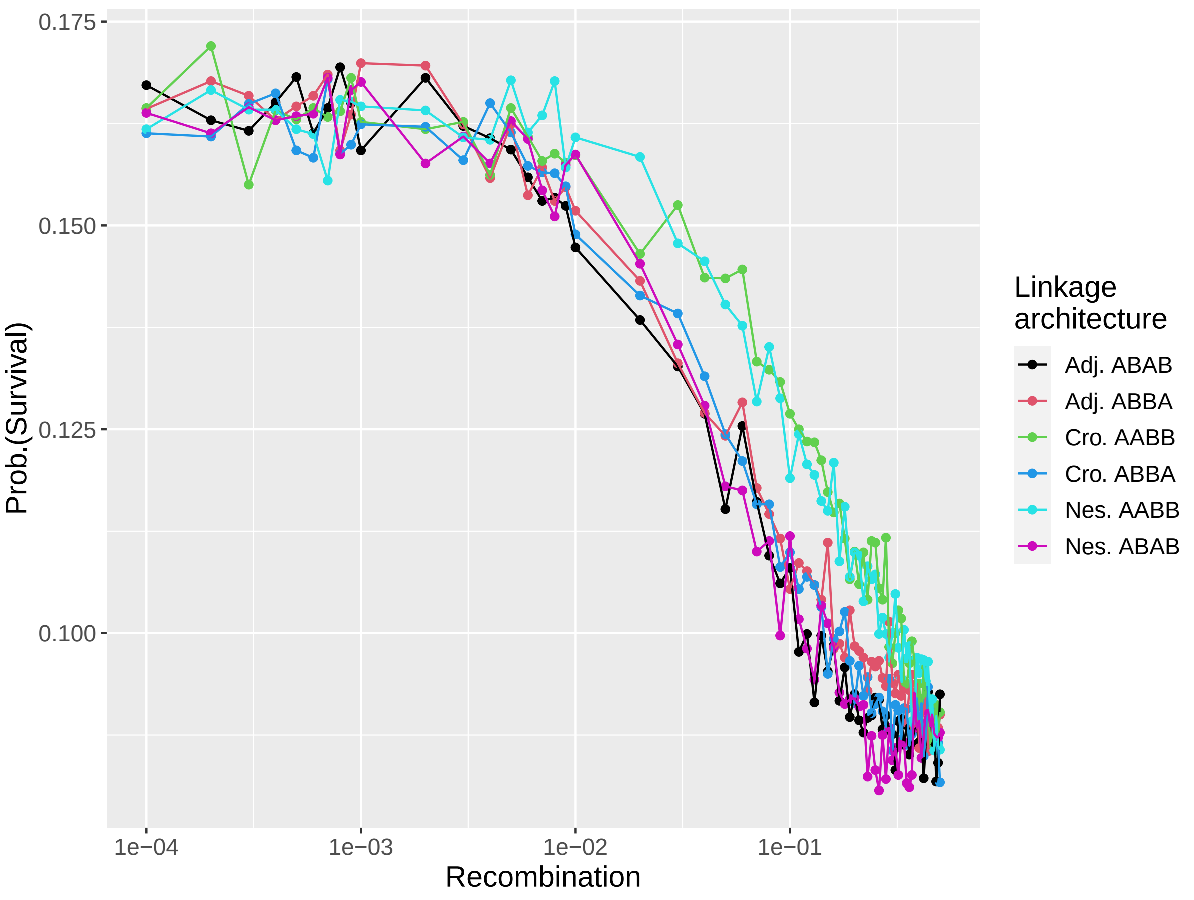


Figure S11: Survival probability as a function of recombination for all 6 architectures for codominant incompatibilities ($\epsilon = -0.2$) and small populations ($N_{\text{init}} = 100$). Additional parameters are set to $\alpha_1 = \alpha_2 = \beta_1 = \beta_2 = -0.001, K = 10^6, n_{\text{rep}} = 10,000$

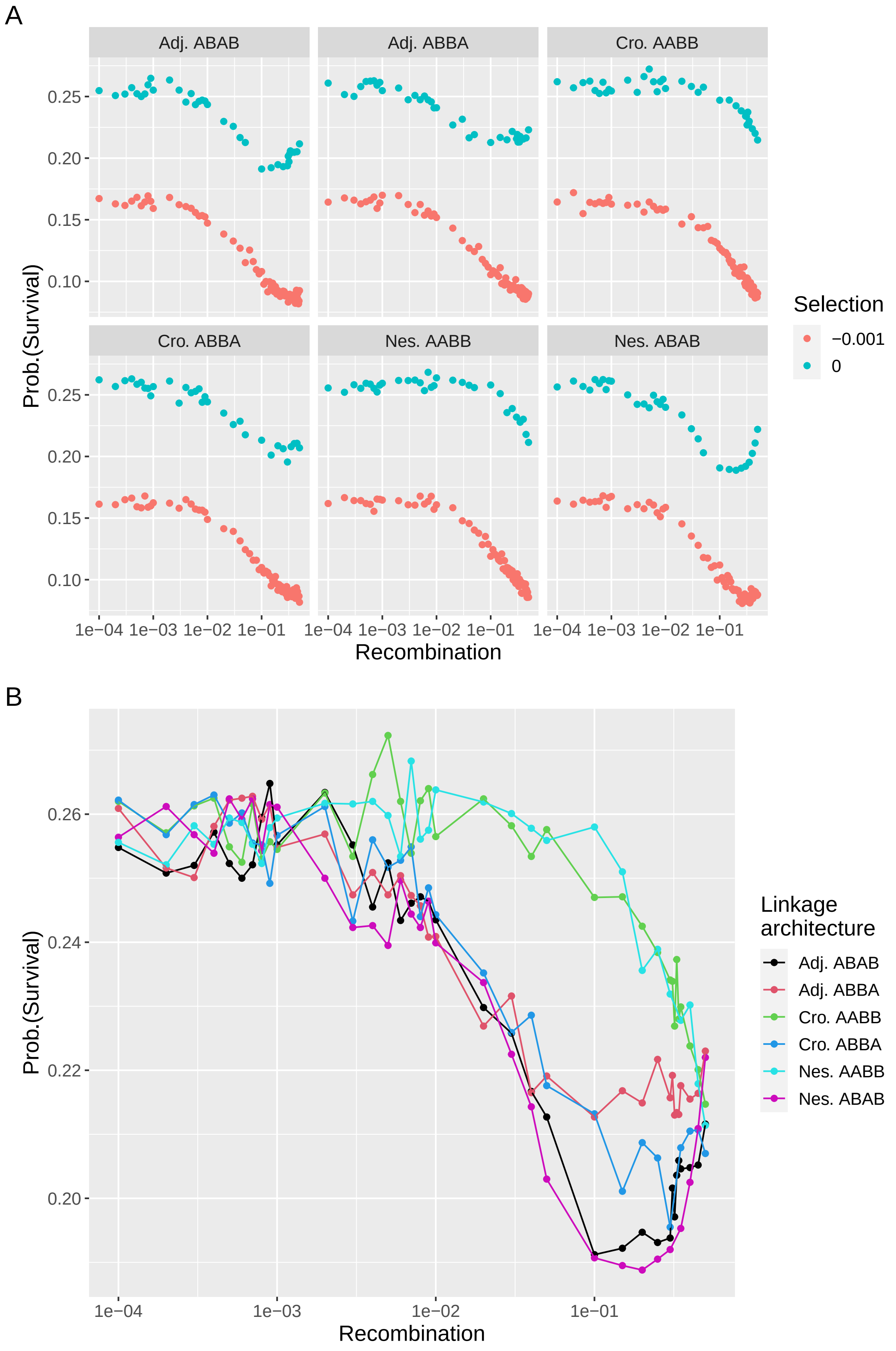


Figure S12: Survival probability of a hybrid population, with an initial population size of $N_{ini} = 100$, and codominant DMIs ($\epsilon = -0.2$). A/ Survival probability in the presence (blue, $\alpha_1 = \alpha_2 = \beta_1 = \beta_2 = -0.001$ and absence (red, $\alpha_1 = \alpha_2 = \beta_1 = \beta_2 = 0$) of selection against the ancestral alleles. B/ Comparison between the 6 different linkage architecture in the absence of direct selection ($\alpha_1 = \alpha_2 = \beta_1 = \beta_2 = 0$). Additional parameters are set to $K = 10^6, n_{\text{rep}} = 10,000$

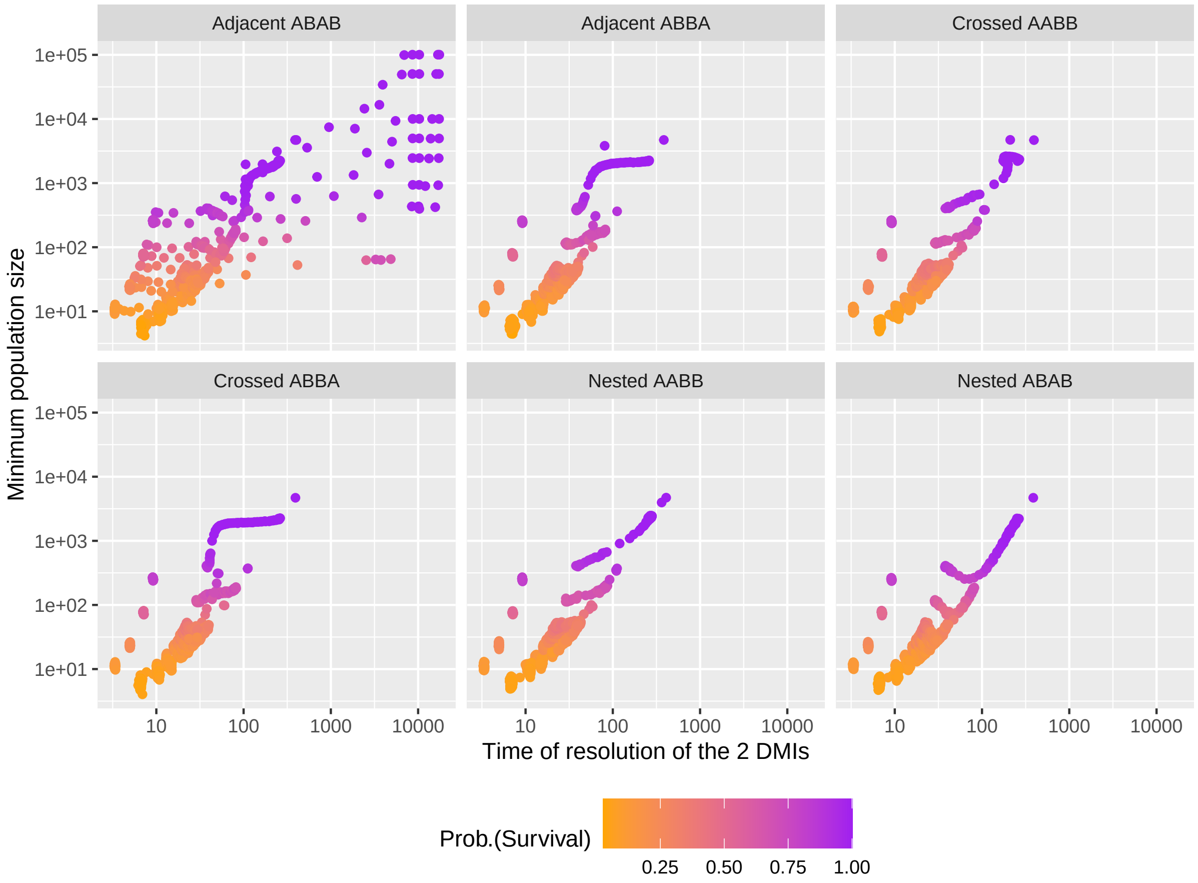


Figure S13: Minimum population size as a function of the resolution time of both DMIs for all 6 linkage architectures. Simulations with different strengths of the DMIs, different genetic distances between loci and different initial population size are pooled together. All genetic incompatibilities are codominant. Color indicates the survival probability. Additional parameters are set to $K = 10^6, n_{\text{rep}} = 1,000$

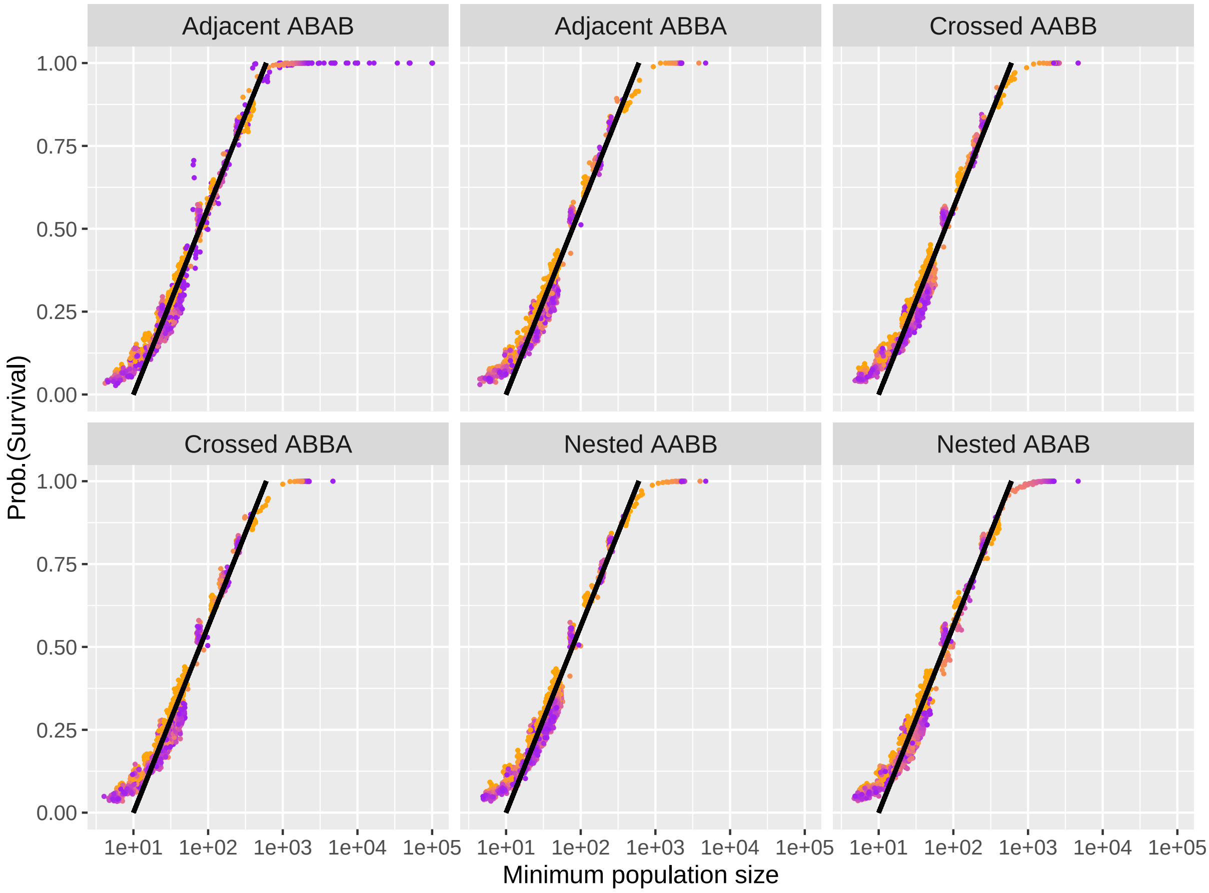


Figure S14: Probability of surviving based on the minimum population size reached during the sorting of the (recessive) genetic incompatibilities for all 6 linkage architectures, for various initial population sizes, recombination rates and epistasis strength. The black line is a visual guideline ($y = -1/59 + x/590$). Color indicates the time (log-scaled) of resolution of the genetic incompatibilities. Additional parameters are set to $K = 10^6, n_{\text{rep}} = 1,000$

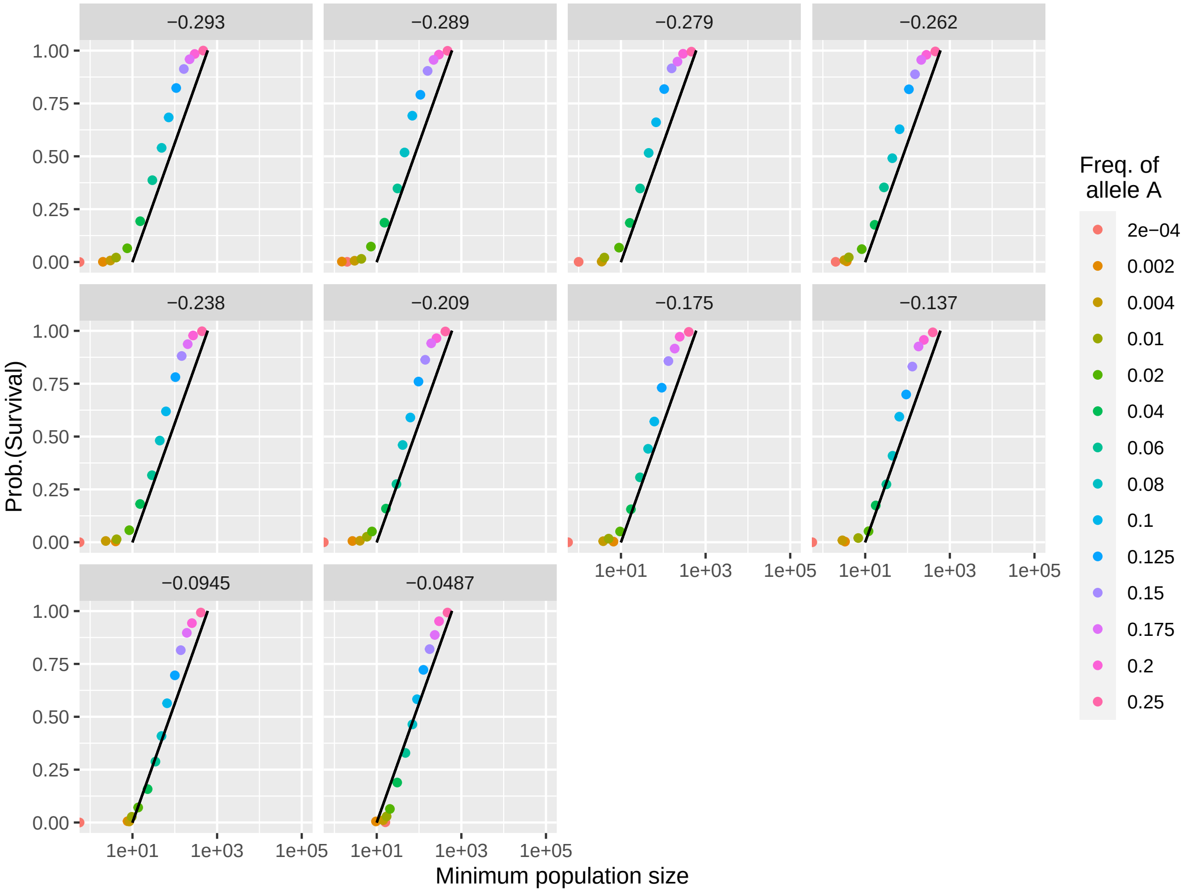


Figure S15: Survival probability of a population as a function of its minimum population size. This figure is the evolutionary rescue scenario equivalent to Figure 5, using the same guideline (y=-1/59+x/590). Different panels correspond to different selective coefficients against allele $a$, such that they generate the same $\bar{r}$ in the initial population (before the introduction of allele $A$) as the F1 hybrid population (see Figure S16). Color indicates the frequency at which allele $A$ is introduced. Additional parameters are set to $K = 10^6, n_{\text{rep}} = 1,000$

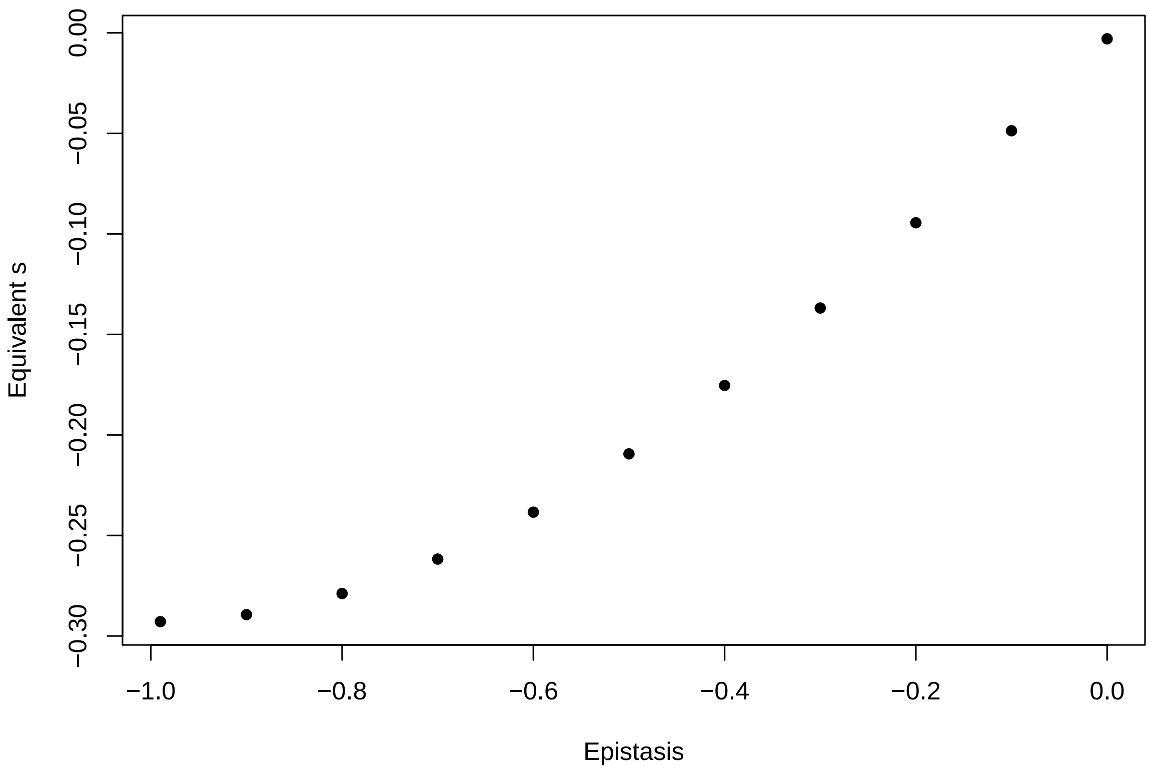


Figure S16: Selection coefficients of the ancestral allele $a$ generating the same initial and final mean growth rate $\bar{r}$ in an evolutionary rescue scenario as the hybrid population affected by codominant genetic incompatibilities. The initial growth rate corresponds to the growth rate of an F1 hybrid individual.

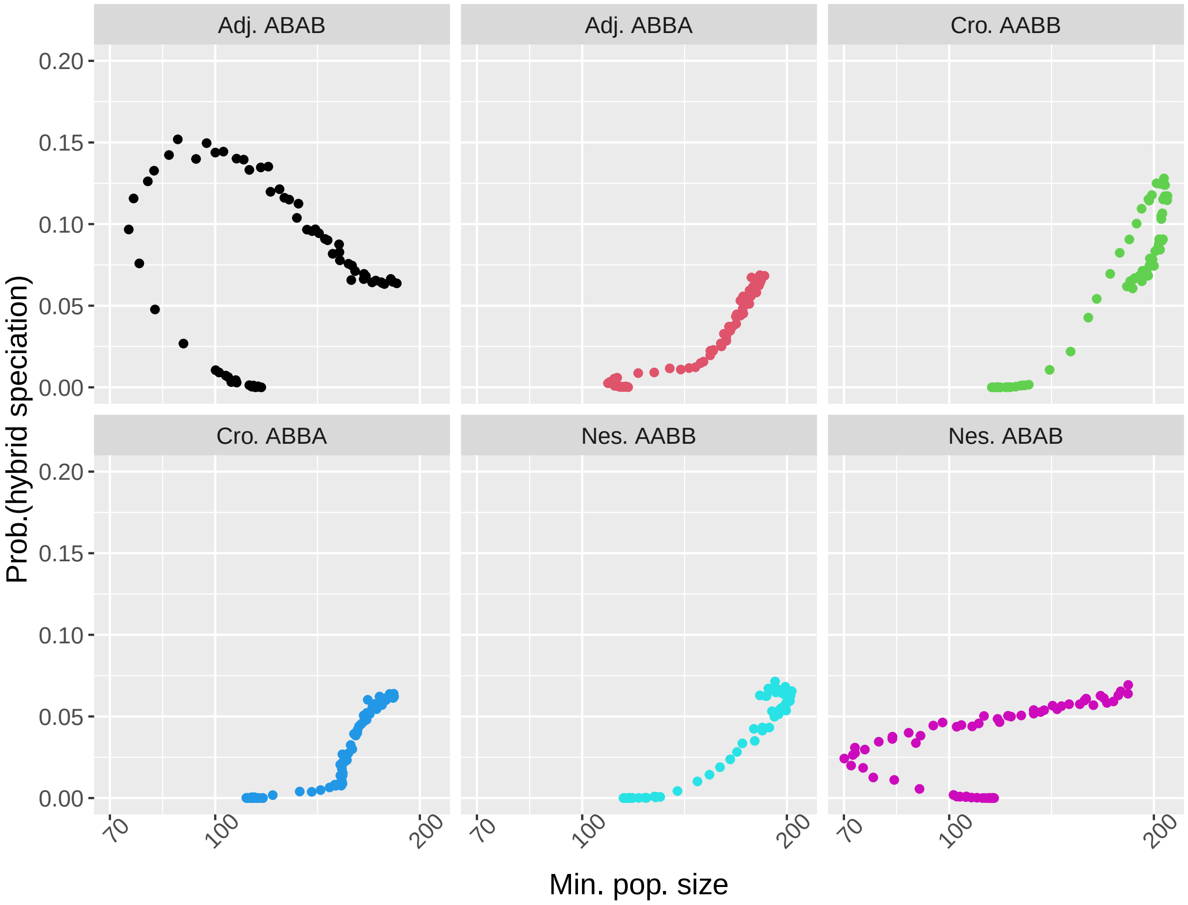


Figure S17: Probability of hybrid speciation is a multivalued function of the minimum population size. Here we considered two pairs of codominant DMIs ($\epsilon = -0.2$) for various genetic distances between loci. This corresponds to the same data that are displayed in Figure 5A in the main manuscript Additional parameters are set to $\alpha_1 = \alpha_2 = \beta_1 = \beta_2 = -0.001, K = 10^6, n_{\text{rep}} = 10,000$

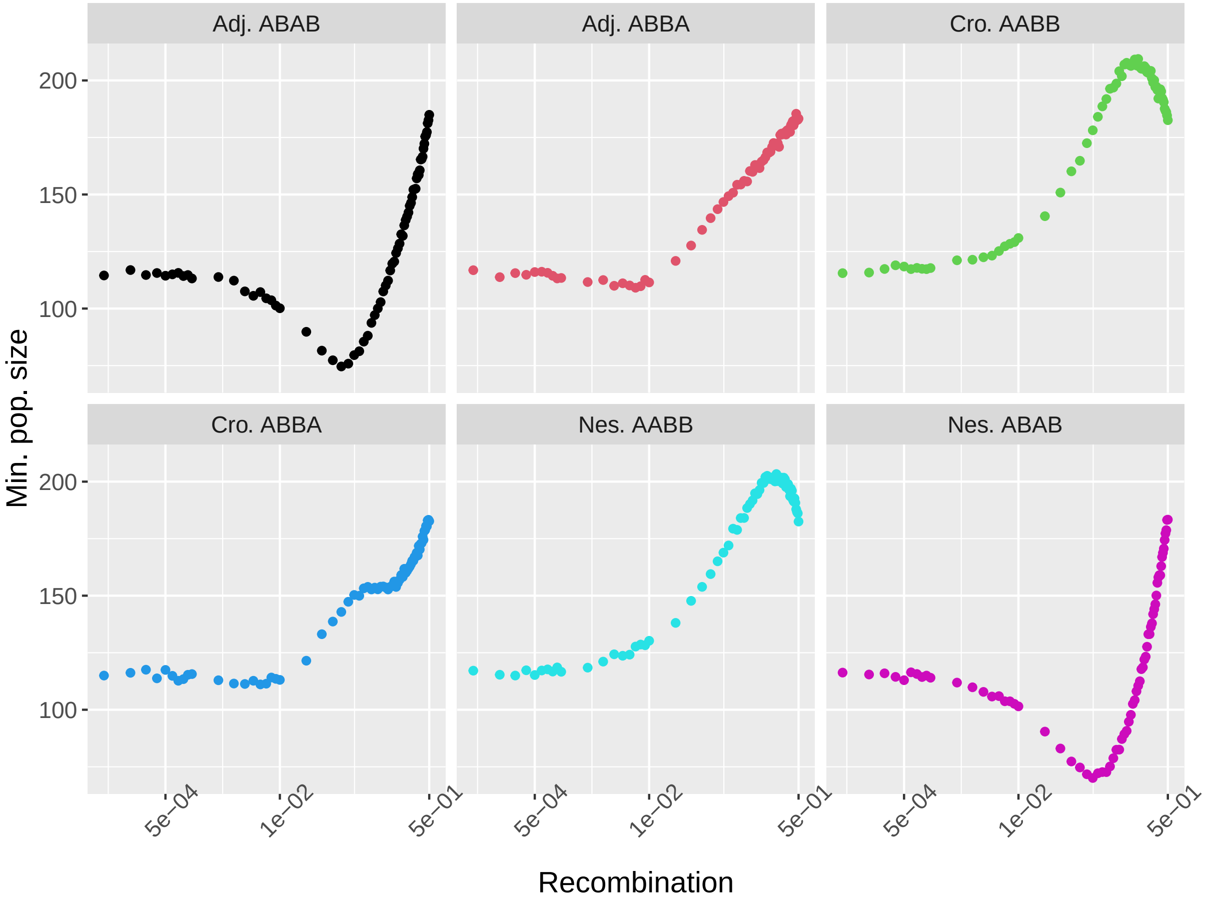


Figure S18: Minimum population size is a non-monotonic function of the genetic distance between loci. Here we considered two pairs of codominant DMIs ($\epsilon = -0.2$). This corresponds to Figure 5A in the main manuscript and in figure S17. Additional parameters are set to $\alpha_1 = \alpha_2 = \beta_1 = \beta_2 = -0.001, K = 10^6, n_{\text{rep}} = 10,000$

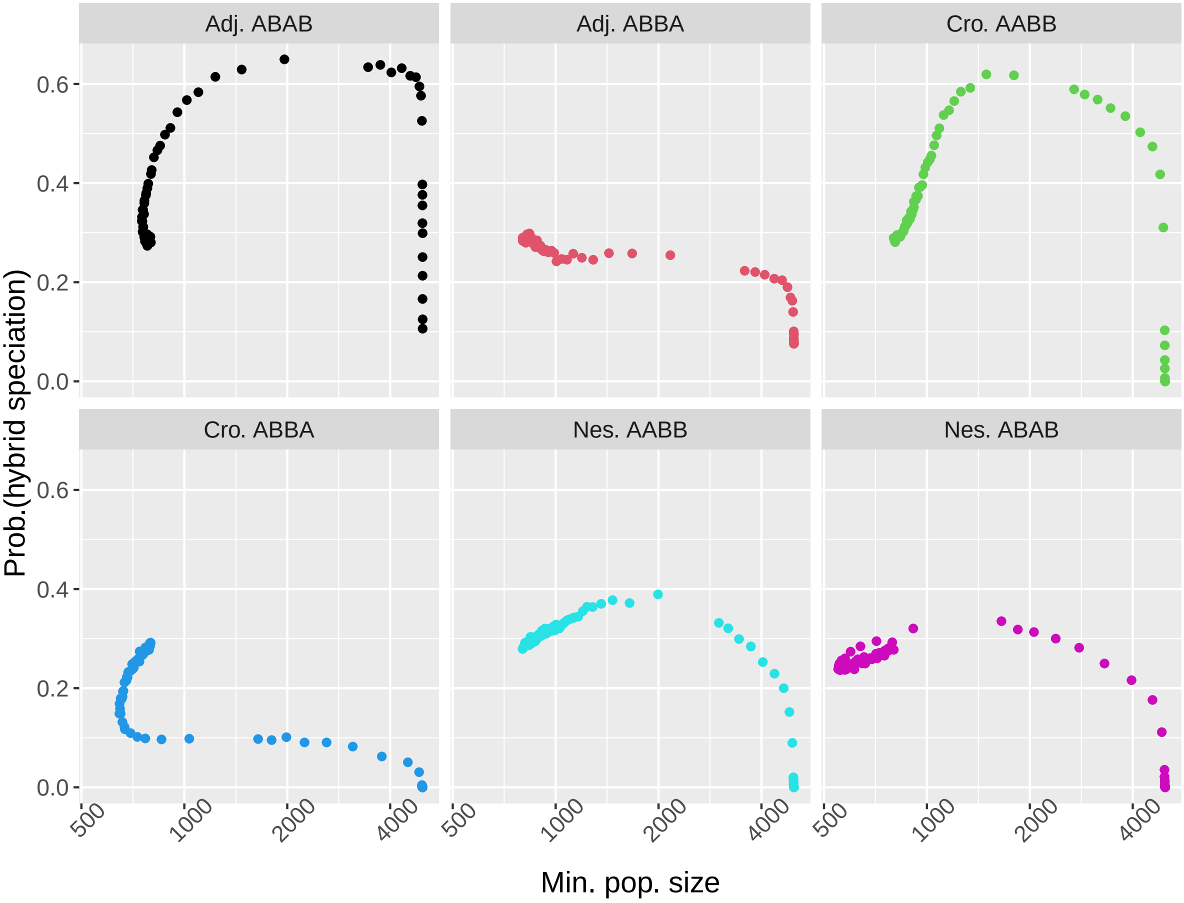


Figure S19: Probability of hybrid speciation is a multivalued function of the minimum population size. Here we considered two pairs of recessive DMIs ($\epsilon = -0.2$) for various genetic distances between loci. This corresponds to Figure 5B in the main manuscript. Additional parameters are set to $\alpha_1 = \alpha_2 = \beta_1 = \beta_2 = -0.001, K = 10^6, n_{\text{rep}} = 10,000$

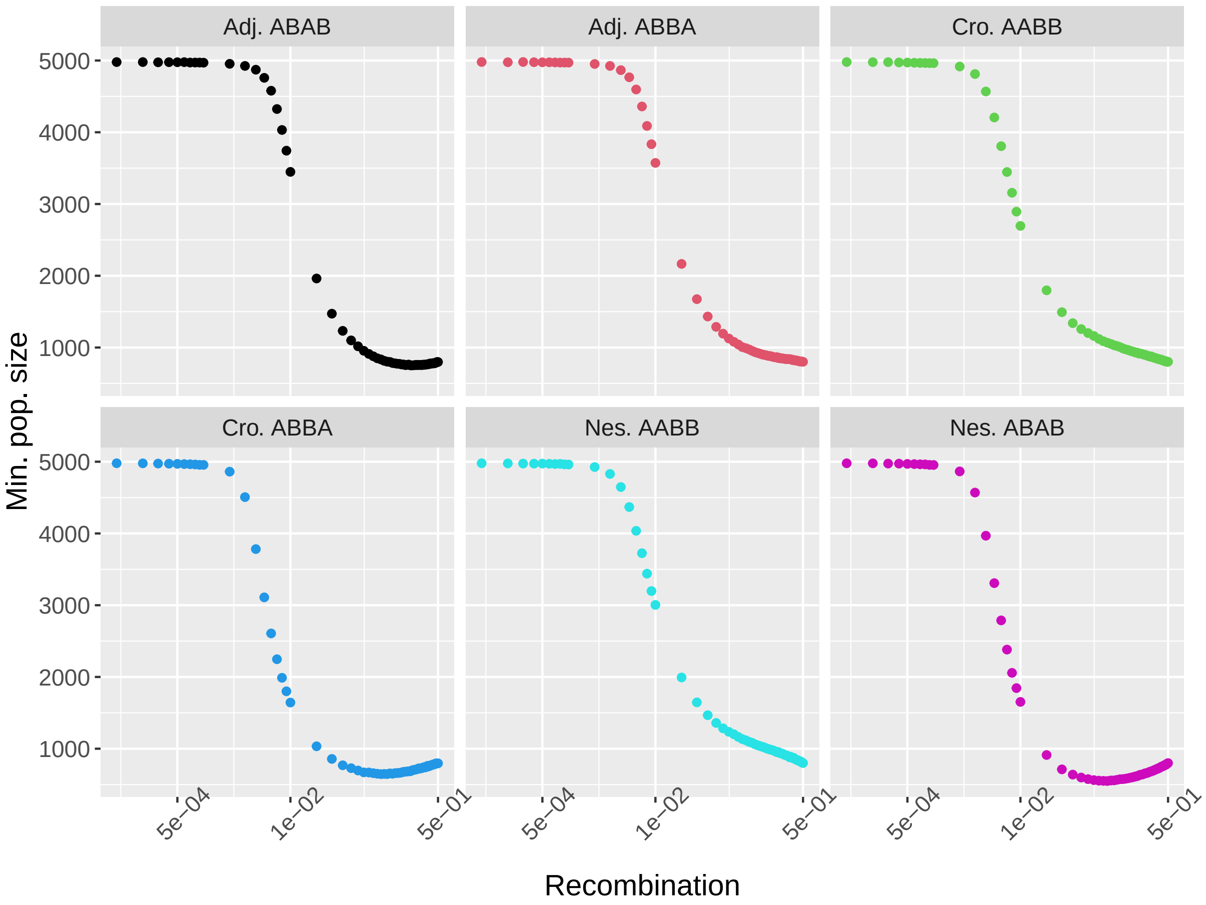


Figure S20: Minimum population size is a non monotonic function of the genetic distance between loci. Here we considered two pairs of recessive DMIs ($\epsilon = 0.2$). This corresponds to Figure 5B in the main manuscript and in figure S19. Additional parameters are set to $\alpha_1 = \alpha_2 = \beta_1 = \beta_2 = -0.001, K = 10^6, n_{\text{rep}} = 10,000$

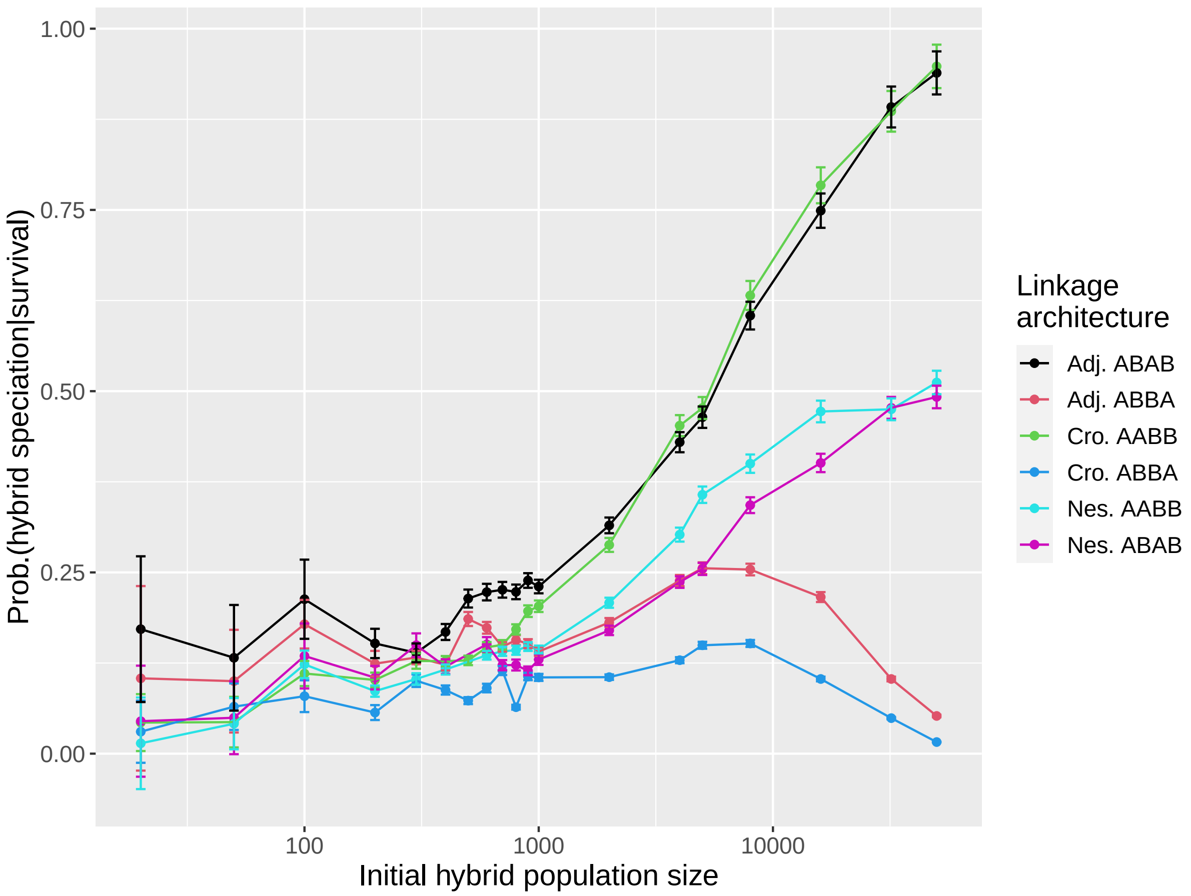


Figure S21: Probability of hybrid speciation, conditioned on survival, as a function of the initial population size for all linkage architecture for two pairs of recessive DMI ($\epsilon = -0.2$). Lines correspond to a local smoothing function and serve as a guide for the eye. It is the recessive equivalent of Figure 6 in the main manuscript. Additional parameters are set to $\alpha_1 = \alpha_2 = \beta_1 = \beta_2 = -0.001, K = 10^6, n_{\text{rep}} = 1,000$

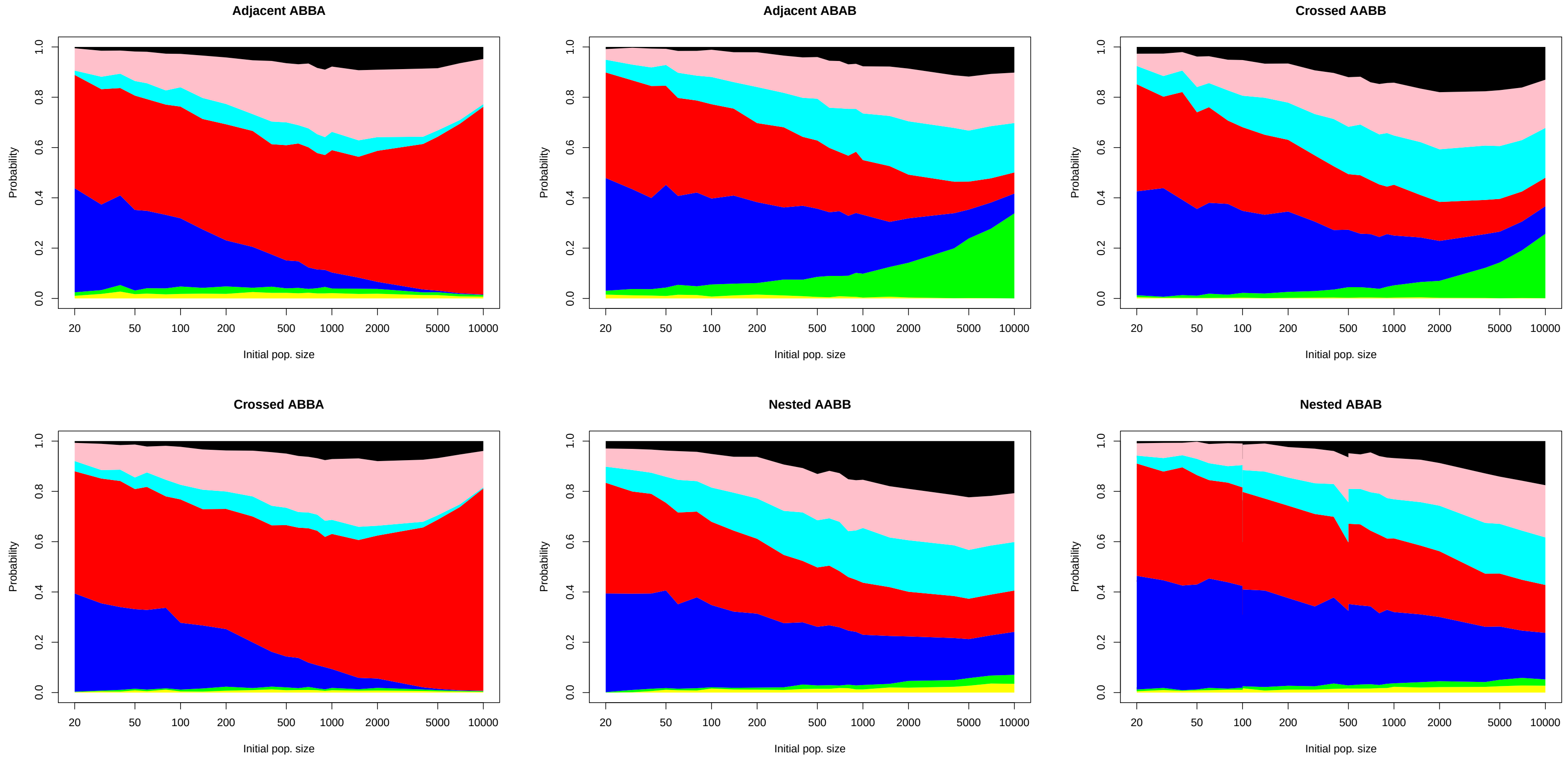


Figure S22: Probability of fixing the different haplotypes for 6 linkage architectures conditioned on survival. Color corresponds to the different genotypes: yellow and green for hybrid speciation haplotypes are given in yellow ($A_1a_2b_1B_2$) and green ($a_1A_2B_1b_2$), red and blue for the parental haplotypes in red ($A_1A_2b_1b_2$) and blue ($a_1a_2B_1B_2$), the haplotypes with one single derived alleles in pink ($A_1a_2b_1b_2$ and $a_1A_2b_1b_2$) and cyan ($a_1a_2B_1b_2$ and $a_1a_2b_1B_2$) and black for the ancestral haplotype ($a_1a_2b_1b_2$). The incompatibilities are codominant. Additional parameters are set to $\alpha_1 = \alpha_2 = \beta_1 = \beta_2 = -0.001, \epsilon = -0.2, K = 10^6, n_{\text{rep}} = 10,000$

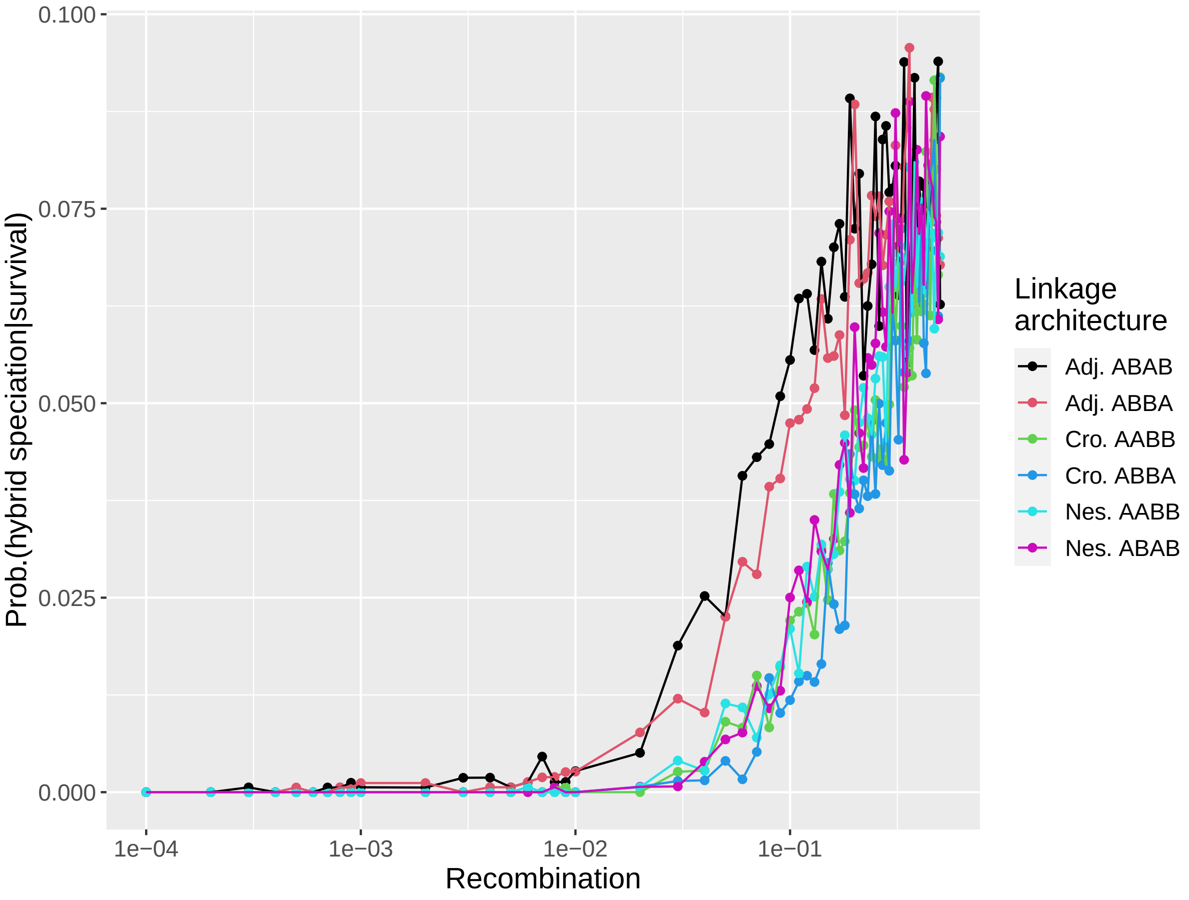


Figure S23: Probability of hybrid speciation, conditioned on survival, as a function of the genetic distance between loci for all linkage architecture for codominant DMIs ($\epsilon = -0.2$) in small population ($N_{\text{init}} = 100$). Additional parameters are set to $\alpha_1 = \alpha_2 = \beta_1 = \beta_2 = -0.001, K = 10^6, n_{\text{rep}} = 10,000$

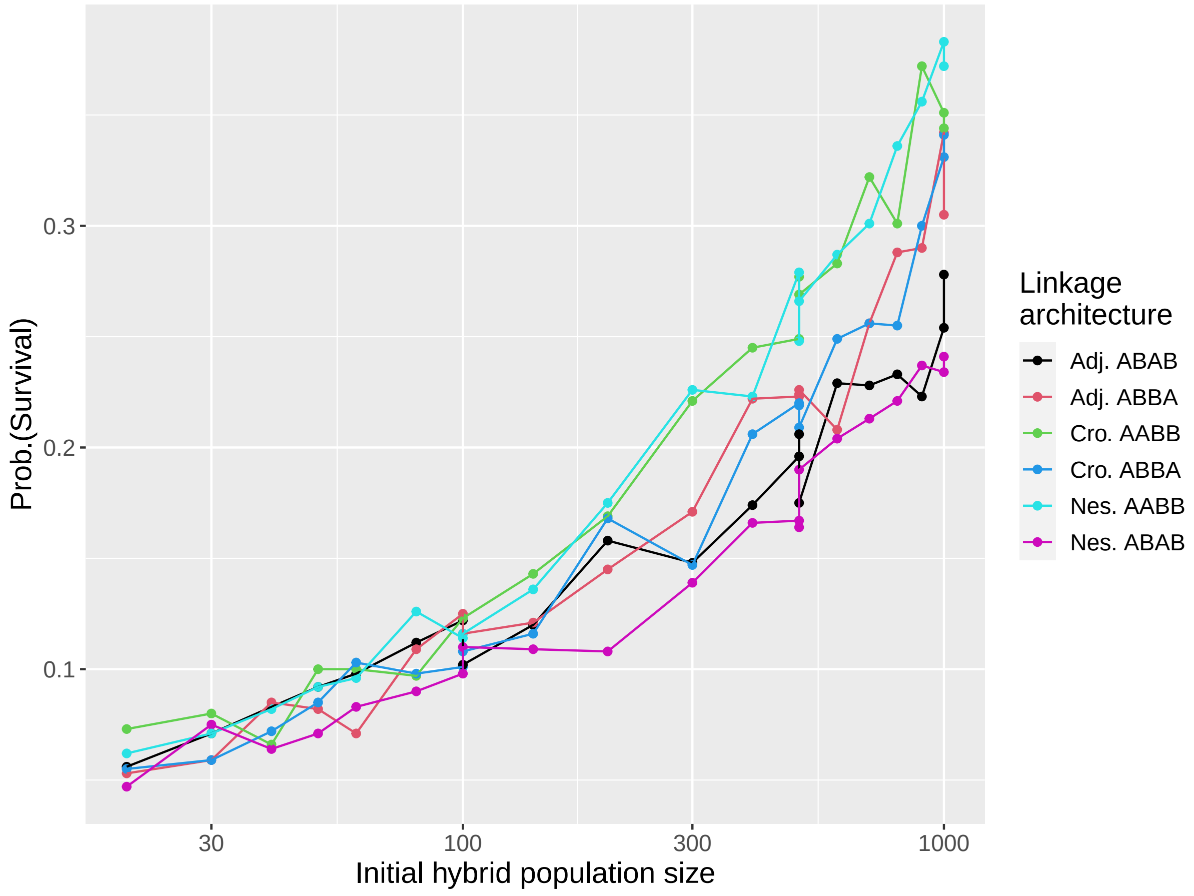


Figure S24: Survival probability for the codominant case ($\epsilon = -0.2$) for all 6 linkage architectures. This corresponds to Figure 6 in the main manuscript. Additional parameters are set to $\alpha_1 = \alpha_2 = \beta_1 = \beta_2 = -0.001, r = 0.1, K = 10^6, n_{\text{rep}} = 1,000$

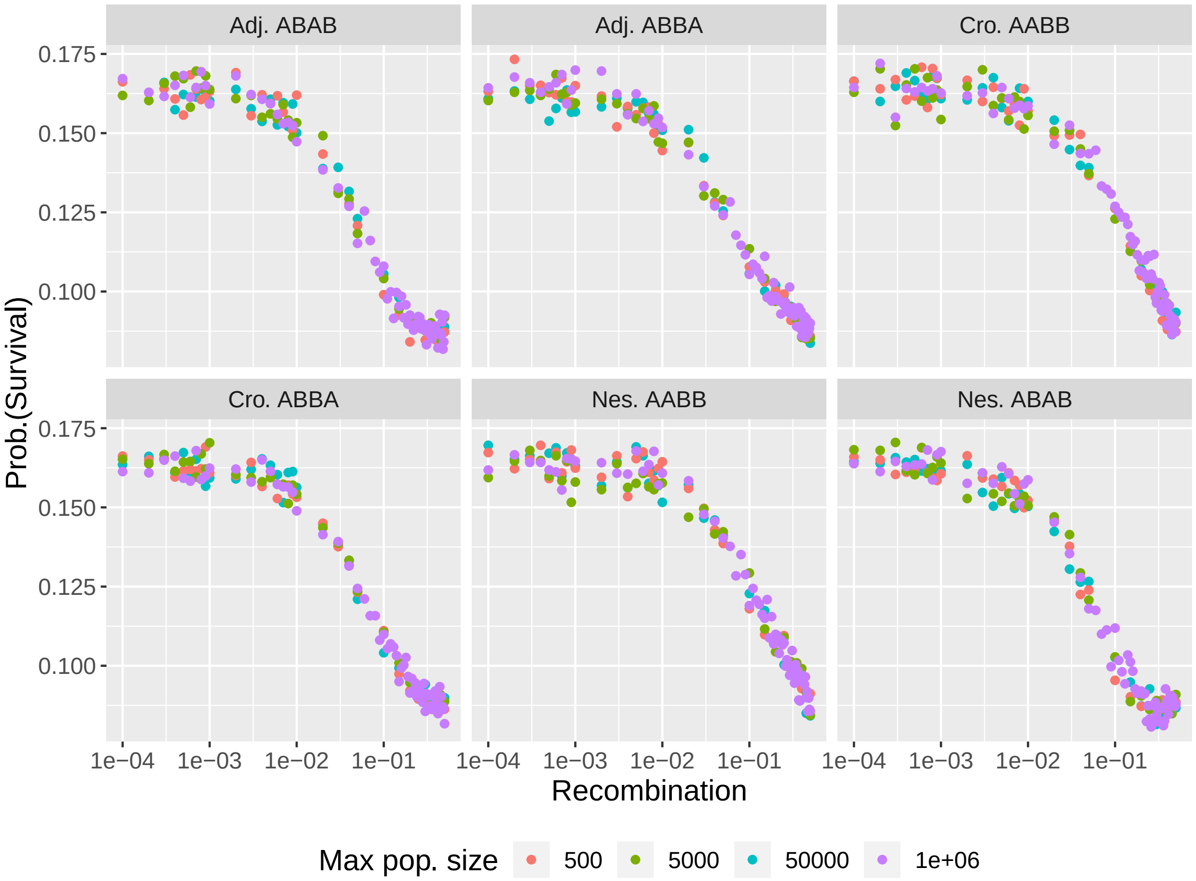


Figure S25: Survival probability does not depend on the carrying capacity of the environment. The incompatibilities are codominant. Additional parameters are set to $N_{\text{init}} = 100, \alpha_1 = \alpha_2 = \beta_1 = \beta_2 = -0.001, \epsilon = -0.2, n_{\text{rep}} = 10,000$

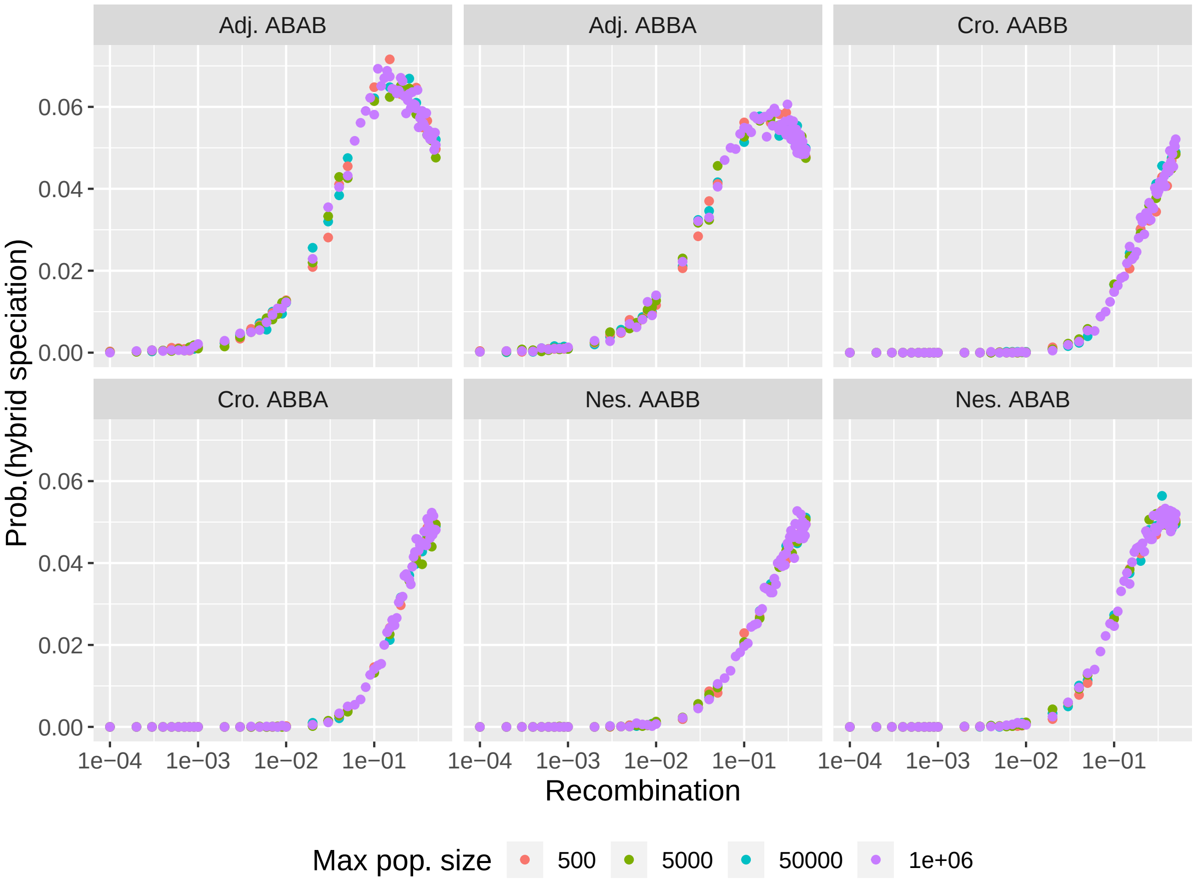


Figure S26: Hybrid speciation probability does not depend on the carrying capacity of the environment. The incompatibilities are codominant. Additional parameters are set to $N_{\text{init}} = 100, \alpha_1 = \alpha_2 = \beta_1 = \beta_2 = -0.001, \epsilon = -0.2, n_{\text{rep}} = 10,000$

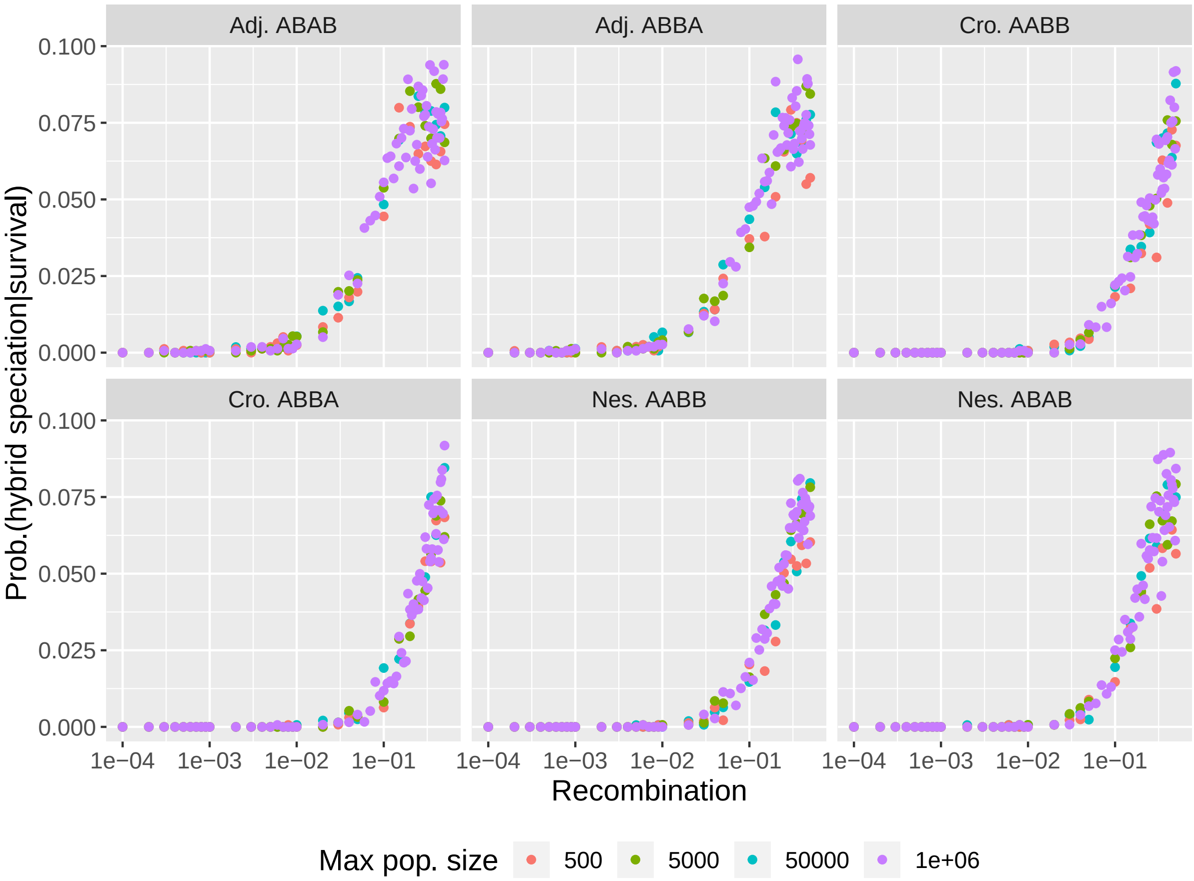


Figure S27: Hybrid speciation probability, conditioned on survival, does not depend on the carrying capacity of the environment. The incompatibilities are codominant. Additional parameters are set to $N_{\text{init}} = 100, \alpha_1 = \alpha_2 = \beta_1 = \beta_2 = -0.001, \epsilon = -0.2, n_{\text{rep}} = 10,000$

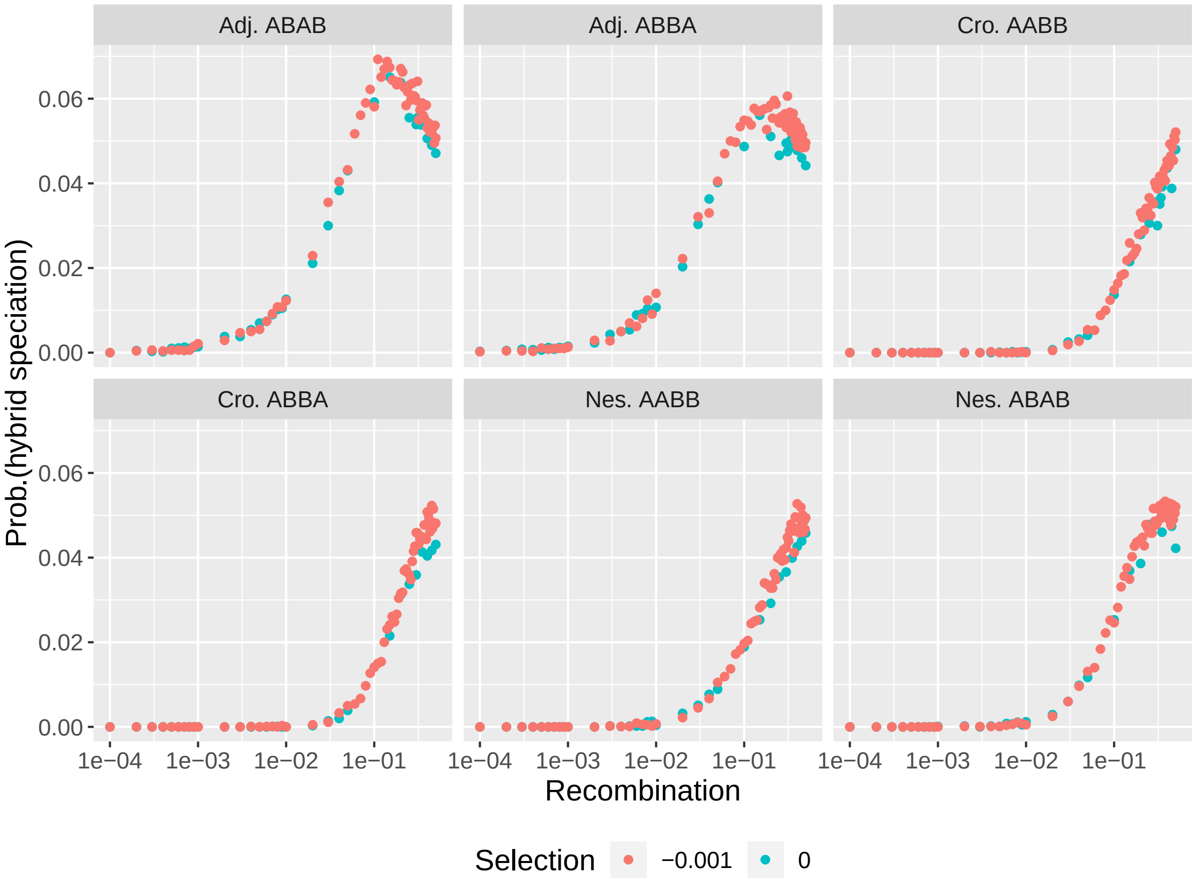


Figure S28: Hybrid speciation probability with and without selection against the ancestral alleles. The incompatibilities are codominant. Additional parameters are set to $N_{\text{init}} = 100, \alpha_1 = \alpha_2 = \beta_1 = \beta_2 = -0.001, \epsilon = -0.2, n_{\text{rep}} = 10,000$

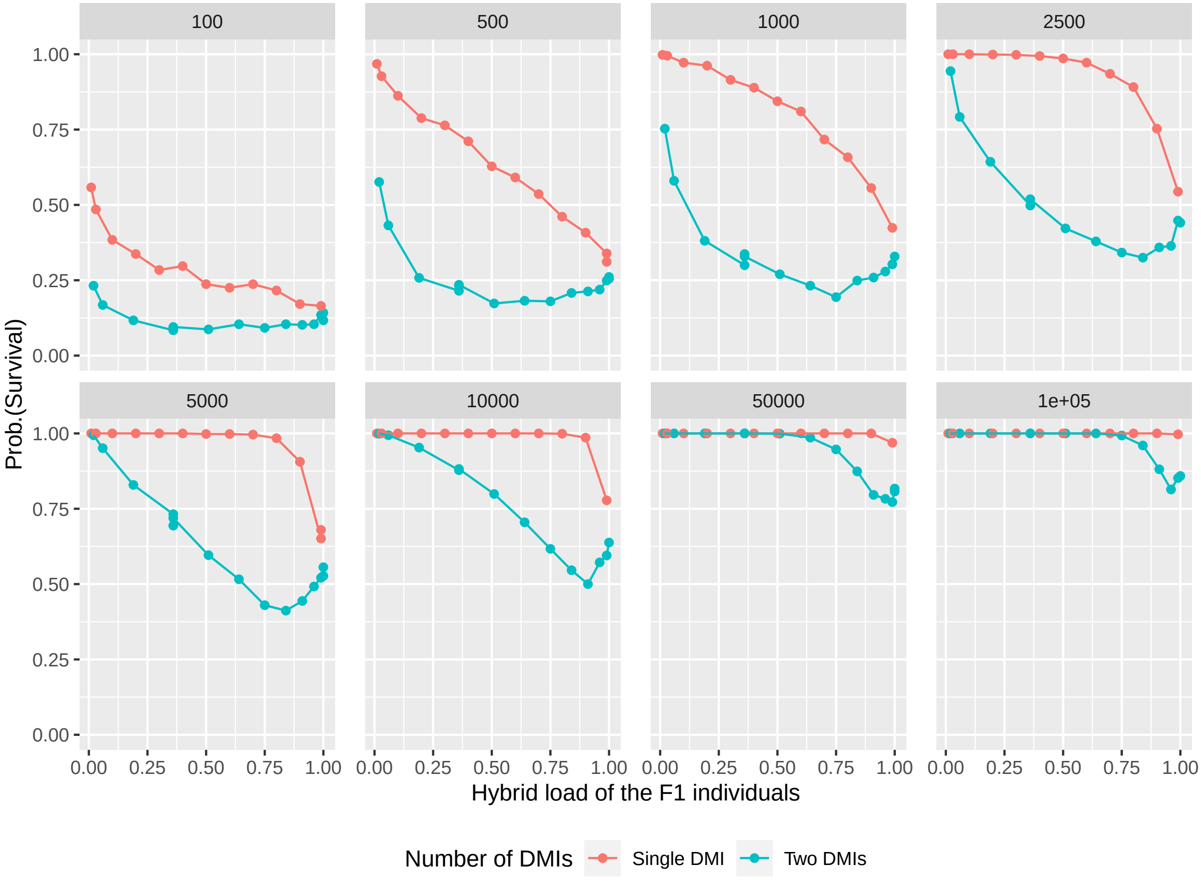


Figure S29: Survival probability as a function of the hybrid load of an F1 individual for codominant unlinked DMIs ($r = 0.5$) for one (red) and two (blue) pairs of DMIs and for different population sizes (given above each panel). Additional parameters are set to $\alpha_1 = \alpha_2 = \beta_1 = \beta_2 = -0.001, n_{\text{rep}} = 1,000$
